# Building Fluorescence Lifetime Maps Photon-by-photon by Leveraging Spatial Correlations

**DOI:** 10.1101/2022.11.29.518311

**Authors:** Mohamadreza Fazel, Sina Jazani, Lorenzo Scipioni, Alexander Vallmitjana, Songning Zhu, Enrico Gratton, Michelle A. Digman, Steve Pressé

## Abstract

Fluorescence lifetime imaging microscopy (FLIM) has become a standard tool in the quantitative analysis of sub-cellular environments. However, quantitative FLIM analyses face several challenges. First, spatial correlations between pixels are often ignored as signal from individual pixels is analyzed independently thereby limiting spatial resolution. Second, existing methods deduce photon ratios instead of absolute lifetime maps. Next, the number of lifetime components contributing to the signal is unknown, while excited state lifetimes with <1 ns difference are difficult to discriminate. Finally, existing analyses require high photon budgets, and often cannot rigorously propagate experimental uncertainty into values over lifetime maps and number of components involved. To overcome all of these challenges simultaneously and self-consistently at once, we propose the first doubly nonparametric framework. That is, we learn the number of fluorescent species (through beta-Bernoulli process priors) and absolute lifetime maps of these species (through Gaussian process priors) by leveraging information from pulses not leading to observed photon. We benchmark our algorithm using a broad range of synthetic and experimental data and demonstrate its robustness across a number of scenarios including cases where we recover lifetime differences between components as small as 0.3 ns with merely 1000 photons.

---

Among many fluorescence microscopy techniques^1–6^, fluorescence lifetime imaging microscopy^1,2^ (FLIM) has been pivotal in probing details and interactions within sub-cellular environments, fluids and solid materials^7–26^. For example, FLIM has been employed in deducing nanoscale maps of optical^8,10–12^, thermodynamic^13–20^, and chemical parameters^21–25^. Furthermore, FLIM is relevant to the health sciences^27–29^ where its sensitivity has been critical in detecting lifetime differences between protonated and unprotonated NAD, an important marker for cancer metabolism^24,30^. What is more, FLIM has also been employed in the field of drug discovery to monitor drug activity within complex biological environments^31,32^.

In typical FLIM experiments, data consist of a series of photon arrival times, following laser pulses, whose statistics are dictated by the present number of fluorescing species and excited state lifetime. Photon arrival times can then be decoded to learn the number of species (or, alternatively, the number of lifetime components) as well as their associated lifetimes. In imaging across regions of space, we may also decode the corresponding lifetime maps.

Here, we assume that the input arrival times (the FLIM data) are collected using a scanning confocal setup. In this setup, a pulsed laser, often with a Gaussian waist, scans the sample at a constant speed over uniformly spaced horizontal trajectories where the spacing defines the pixel size. The excited fluorophores then emit photons with a random delay drawn from a distribution characteristic of the fluorophore species, Fig. 1a. Moreover, the recorded arrival times are further contaminated by instrumental noise. That is: 1) the detector delay in recording the arrival times; and 2) the unknown exact time of excitation due to the finite breadth of excitation pulses. Together, these are often modeled using a Gaussian distribution, termed the instrumental response function (IRF). The exponential waiting time for de-excitation from each lifetime component and the effects of the IRF thus result in two layers of stochasticity in reported photon arrival times given by the convolution of the exponential and IRF distributions.

**Figure 1:**
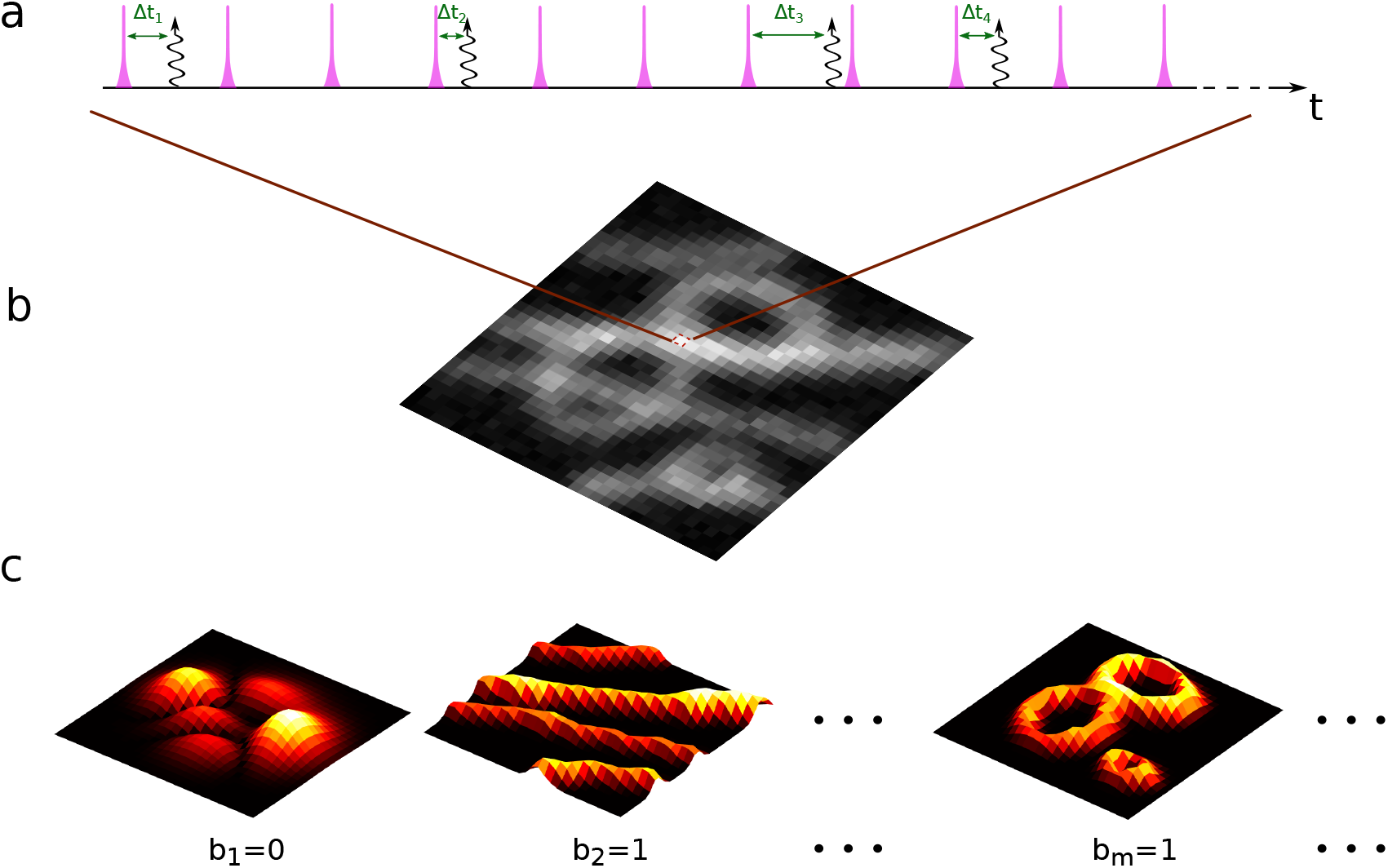
A cartoon illustration of a typical FLIM experiment and how the BNP-FLIM algorithm works. (a) Every spot in the specimen is illuminated by a train of laser pulses, designated by pink spikes, where a fraction of them lead to the detection of photons, shown by curly arrows. The photon arrival times, Δ*t_k_*, are recorded and used in FLIM analysis to infer the number of lifetime components as well as their associated spatial maps and corresponding lifetimes. (b) The sets of photons drawn from all spots is arranged into a two dimensional pixel array representing the raw FLIM data. (c) The Bayesian nonparametric FLIM (BNP-FLIM) algorithm models the input data (nominally) assuming an infinite number of lifetime components. To each lifetime component is associated an nominally infinite number of candidate spatial maps for how the component is distributed. Eventually, as shown in (c), our method determines: 1) which lifetime component are warranted by the data (for which the associated Bernoulli variable, *b*, is found to be unity) and what its lifetime is; and 2) its associated lifetime map. In the case shown in (c), only the second and *m*^th^ components are warranted by the data and have a non-zero associated Bernoulli variable b (*i.e., b*_2_ = *b_m_* = 1). The map determined for the first component (with *b*_1_ = 0) is thus immaterial.

To learn the number of lifetime components as well as their associated lifetimes from FLIM data, the community relies either on model-free methods, such as phasor-based approaches^33–36^ and neural networks^37–39^, or model-based methods such as FLIM data analysis techniques. For instance, model-based techniques include: time correlated single photon counting (TCSPC) histogram fitting to extract lifetimes from photon arrival histograms via least square fitting^40–42^; photon-by-photon likelihood maximization^43–45^; construction of Bayesian posteriors over lifetimes warranted by the data^46–52^; and classic deconvolution methods ^53–55^.

In spite of this progress in FLIM data analysis, only one such technique does not assume a number of lifetime components *a priori* while propagating unavoidable error induced by the stochastic processes above over those numbers of components that may be warranted by the data^56^. Even so, this technique is limited to a single pixel (*i.e*., single illuminated confocal spot) and therefore cannot leverage spatial correlations across pixels to extract high resolution lifetime maps required to smoothly and quantitatively interpolate lifetime maps between and below pixel areas.

While heuristics exist to deduce the number of lifetime components^57^, these do not propagate error and rely on data pre-processing which fundamentally limits the ability of these methods to separate close lifetimes. Indeed, other existing techniques, save the one above, require prior knowledge of the number of lifetime components^37–44,46–50,58^ while, when analyzing images, analyze them in a spot-by-spot spatially decorrelated manner^33,34,42,47–49,56^.

The above motivates the reason why it is important to avoid data pre-processing (such as histogram fitting by TCSPC) in order to learn the number of components (as opposed to specifying them by hand) while simultaneously deducing their spatial lifetime maps. To be clear, even hypothesized single lifetime components may further split into multiple components on the basis of the chemical environments to which they are exposed within a cell^59,60^. What is more, these components may have similar lifetimes and exist in spatially overlapping regions further highlighting the importance of propagating error and accounting for spatial correlation across pixels while satisfying the underlying Poisson emission statistics.

To compensate for the loss of information in data pre-processing, multiple methods therefore require large photon budgets that otherwise carry the risk of specimen photo-damage^38,42,49^. Here, by contrast, our objective is to simultaneously deduce: 1) the number of lifetime components present within a given FLIM dataset; while 2) learning absolute lifetime maps with sub-nanosecond temporal resolution, *i.e*., distinguishing lifetimes with sub-nanosecond difference, and interpolating lifetime maps below sub-pixel for each lifetime component determined in the first point. We do so by leveraging the spatial correlations between pixels and information within pulses (termed empty pulses) that do not lead to any photon observation. We develop a framework within the Bayesian paradigm precisely to propagate all sources of errors over items 1-2 above.

Yet, since the number of lifetime components, and their associated maps are unknown, we must operate within a Bayesian nonparametric framework illustrated in Fig. 1 which is, in fact, doubly nonparametric.

To be precise, we invoke the beta-Bernoulli process prior^61–63^ over each candidate component. That is, we associate a binary weight, *b* ∈ {0,1}, to each map where non-zero weights are ascribed to those lifetime maps warranted by the data. As such, the number of lifetime maps present in a given data set are enumerated as the number of non-zero weights, Fig. 1c. Next, as our goal is to determine what spatial maps for each species determined are warranted by the data of all possible candidate lifetime maps (of a nominally infinite number), we invoke Gaussian process (GP) priors^64–67^. This is critical in allowing us to learn continuous lifetime maps smoothly over large spatial regions^58^, rather than to rely on concatenated pixel-wise maps derived from independent pixel analyses. As we will see, a value for the lifetime map can be deduced at any point in physical space, of the infinitely dense points in the two dimensional focus plane, even below a pixel value with correctly propagated uncertainty.

In what follows, we show that our Bayesian nonparamteric FLIM (BNP-FLIM) algorithm learns the number of lifetime components (with corresponding lifetimes with sub-nanosecond resolution), and interpolating lifetime maps below pixel size using limited FLIM data from experiments described above.

## Methods

Here, we briefly discuss the mathematical framework for our BNP-FLIM algorithm. Full model details developed herein can be found in the Supplementary Information.

BNP-FLIM starts from stochastic photon arrivals, designated by 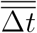, where the double overbar represents the entire set of photons from all pixels. The stochasticity is introduced from the inherent random nature of photon emission and detection process. Stochastic photon arrival times motivate our probabilistic analysis framework.

Our framework starts from the likelihood

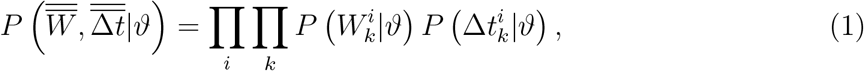

where 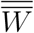 represents a set of binary random variables 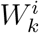 for the *k*th pulse in the *i*th pixel indicating whether the pulse leads to a photon detection or not (empty or non-empty pulse). Also, *ϑ* collects the set of parameters we wish to infer including: lifetime values (*τ_m_*), lifetime maps (Λ_*m*_), the means of GPs (*ν_m_*) on which we place a hyperprior, and the binary weights (*b_m_*) associated to each lifetime component. As we will see, the binary weights are Bernoulli random variables realized to unity for existing lifetime components warranted by the data and 0 otherwise. Here *m* indexes the number of lifetimes and *m* runs from 1: *M*. In Bayesian nonparametrics, we consider *M* → ∞ *a priori* and determine which lifetimes ultimately are warranted by the data.

Furthermore, 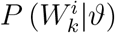 and 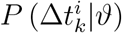, respectively, denote the likelihood of an individual pulse and photon arrival time. In what follows, we describe each of these likelihoods in more detail.

We start by describing the likelihood for the binary observation 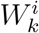. This parameter follows a Bernoulli distribution with success probability of observing a photon of 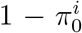 leading to the following likelihood

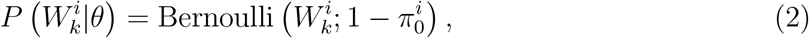

where 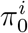 denotes the probability of observing no photons from the *i*th pixel. The explicit form for 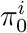 will be derived shortly.

The next observation coincides with the photon arrival times from non-empty pulses, 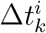, which can originate from any of the species present in input data. To obtain the likelihood of photon arrival times, 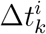, we need to take into account the IRF and sum over all possible events leading to a photon observation, including: all *N* previous pulses that could have excited the fluorophore; and all *M* fluorophore species that might have lead to this photon. This returns

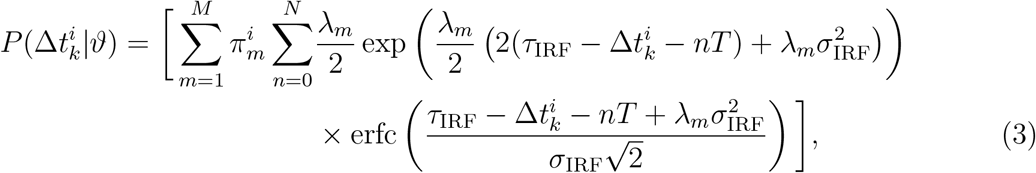

where 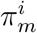 gives the probability of detecting a photon from the *m*th fluorophore species in the *i*th pixel. Here, *λ_m_, τ*_IRF_, *σ*_IRF_ and *T*, respectively, denote the inverse of lifetime (1/*τ_m_*), offset, and variance of the IRF and the inter-pulse time. Derivation details are provided in the Supplementary Information.

After introducing likelihoods in broad terms, we now turn our attention to the probabilities 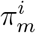 and 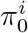 appearing within our likelihoods. These quantities are directly related to the lifetime maps as follows^58^

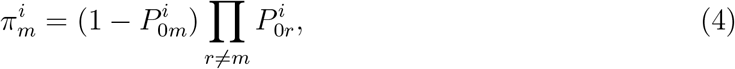

where *P*_0*m*_ denotes the probability of no photon observation from the mth species given by^58^

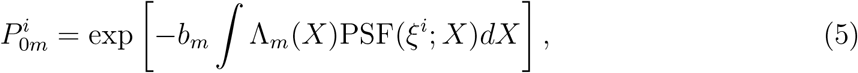

where PSF, *ξ_i_* and *X*, respectively, denote the confocal point spread function, the center of the *i*th pixel corresponding to the center of PSF, and three dimensional spatial coordinates. Furthermore, Λ_*m*_(*X*) is set to *μ_m_*(*X*)*ρ_m_*(*X*) where *μ_m_*(*X*) denotes the excitation probability of a fluorophore of type *m* during a single pulse, and *ρ_m_*(*X*) denotes the concentration of fluorophores^58^. Here, for lifetime maps with zero weights (*b_m_* = 0), eq. 5 reduces to unity leading to zero probability of observing photons from the corresponding species.

Here, we learn absolute lifetime maps by leveraging the information carried by pulses leading to no photon observation, termed empty pulses. This is by contrast to previous FLIM analyses^46–51,56^ that only take into account the set of observed photons from different species in pixel *i*, and learn the ratio of lifetime maps by calculating probabilities of photon observations from different species, 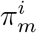.

Here, the probability of observing no photon from the *i*th pixel is given by the product of probabilities of no photon observation from all species

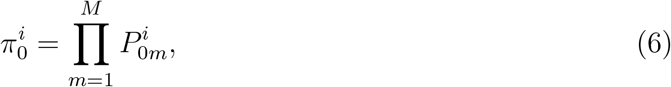

where 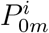 is the probability of no photon detection from species *m*. Considering empty pulses, the sum of these probabilities is naturally unity

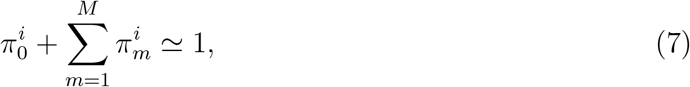

under the assumption that there is no more than a single detected photon per pulse^58^. To build further intuition, removing the contribution from empty pulses 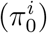 from the above equation results in a sum no longer equal to unity. As such, methods ignoring empty pulses rescale probabilities 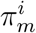 to make them resum to unity and, as a result, only learn rescaled values, *i.e*., photon ratios, rather than the absolute values.

Now, with the likelihood at hand, we proceed to construct the posterior proportional to the product of the likelihood and priors on the unknown parameters

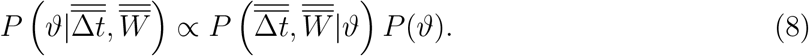

The most notable priors in the BNP-FLIM algorithm are the nonparametric GP priors on the set of lifetime maps and the nonparametric beta-Bernoulli process priors^61–63^ on the binary weights

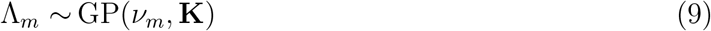

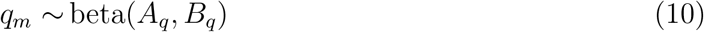

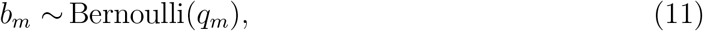

where *ν_m_*, **K** are, respectively, the GP prior mean and the covariance matrix. *A_q_*, *B_q_* are beta-Bernoulli process hyperparameters. The remaining priors are discussed in Supplementary Note 1-2.

As the posterior does not attain an analytic form, it cannot be sampled directly for all variables. Thus the BNP-FLIM algorithm provides a means to sample our posterior to draw inferences about unknown parameters using Marov Chain Monte Carlo techniques ^63,68–75^ (see details in Supplementary Note 2). Furthermore, while nominally the lifetime components are infinite (*M* → ∞) within the nonparametric paradigm, here we instead follow Refs.^61–63^ and set a large upper limit on the number of lifetime components (*M*) in lieu of infinity.

More specifically, our MCMC chain is structured in such a way as to sweep through the entire set of parameters (*i.e*., via Gibbs sampling) at every iteration by independently drawing samples from each parameters’ conditional posterior; see Supplementary Information Note 2. To do so, we sweep the parameter set in the following order:

- lifetime maps, Λ_*m*_–these can be sampled either using the conceptually simpler Metropolis Hastings (MH)^76,77^ or the more efficient elliptical slice sampling^78^. We opt for the latter. Either method of sampling for lifetime maps are required as the likelihood derived in eqs. 1–3 is not conjugate to the GP prior.
- mean of GP priors, *ν_m_*– we use MH to sample the GP means, *ν_m_* due to the lack of conjugacy between the prior and likelihood;
- lifetimes, *τ_m_*–we sample the lifetimes, *τ_m_*, again using MH because again due to lack of conjugacy;
- the binary weights, *b_m_*–we update the binary weights, *b_m_*, by modifying a subset of *b_m_* by randomly picking a subset and directly sampling from the posterior for that subset (See Supplementary Note 2).

In the end, the chain of samples drawn are used for subsequent numerical analysis.

## Results

The overarching aim of our BNP-FLIM algorithm described above is to develop the posterior and sample from it to learn the number of lifetime components, *i.e*., number of non-zero binary weights *b_m_*, warranted by the data, the corresponding lifetimes, *τ_m_*, and lifetime maps, designated by *Λ_1:M_*, using a set of input photon arrival times 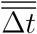 and pulse occupancies 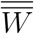. To do so, the BNP-FLIM algorithm draws numerical samples from our posterior for these parameters as illustrated in the former section. Our results are thus histograms over numerical samples drawn from our Monte Carlo sampling scheme. Uncertainties naturally follow from the widths of our histograms reflecting our marginal posteriors.

In this section, we use a range of synthetic and experimental data to benchmark our algorithm against different lifetime maps including: 1) simple, smoothly varying, homogeneous lifetime maps (see Fig. 2 and Fig. S1); 2) more difficult lifetime maps with heterogeneous features similar to those present in experimental samples (see Figs. 3–5 and Figs. S2-7). In addition, employing simulated data with smoothly varying lifetime maps, we further evaluate our algorithm’s performance over a wide range of lifetimes, lifetime differences, and photon counts (see Fig. 2).

**Figure 2:**
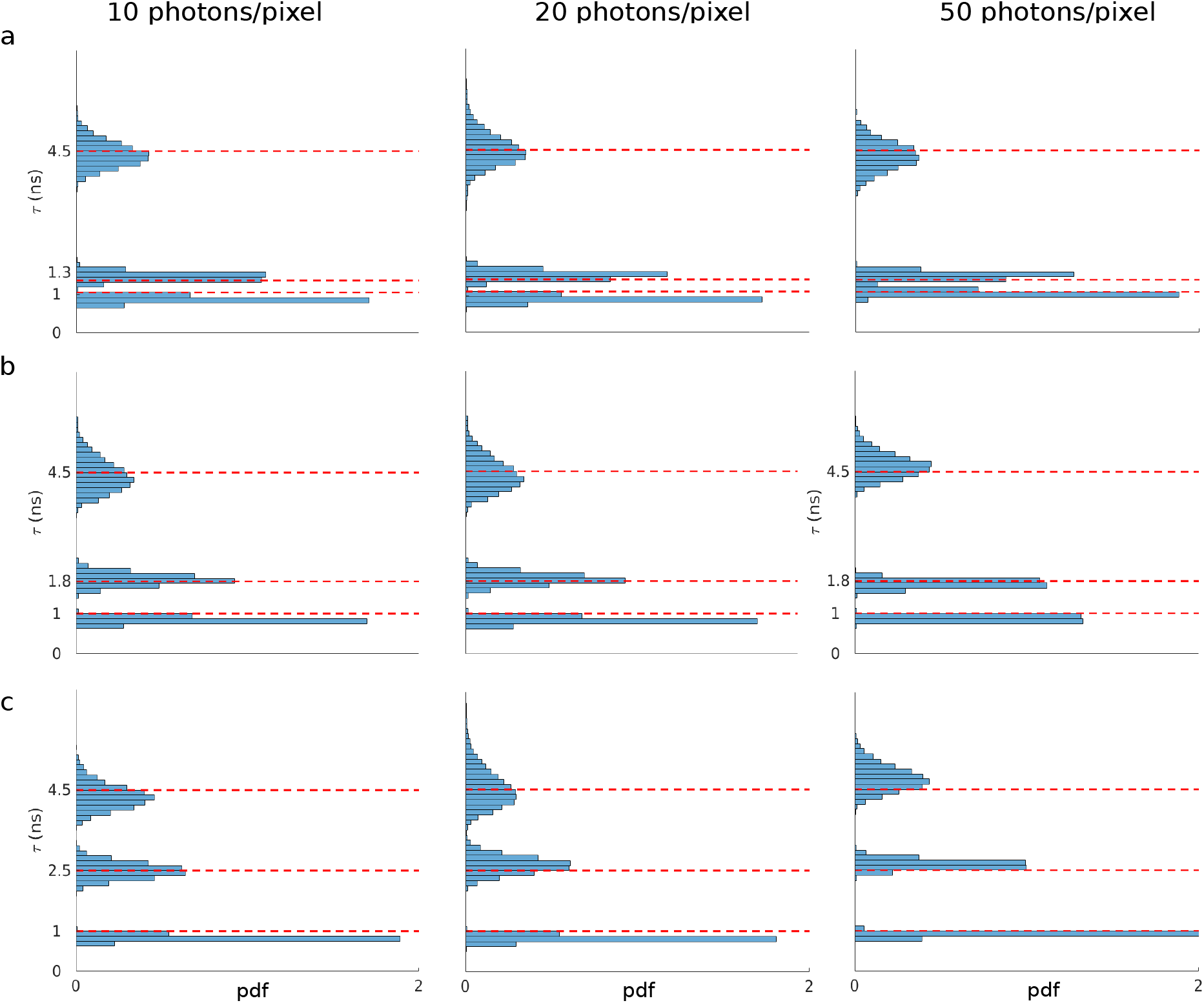
BNP-FLIM robustness with respect to photon counts per pixel and lifetime differences. Three overlapping lifetime maps with different lifetimes were generated over a region of 5×20 pixels and processed by the BNP-FLIM algorithm. There are two fixed lifetimes of 1 ns and 4.5 ns in all simulated data, while we varied the third lifetimes to obtain lifetime differences of (a) 0.3 ns; (b) 0.8 ns and 1.5 ns. Histograms show the resulting lifetime samples from the BNP-FLIM algorithm.

**Figure 3:**
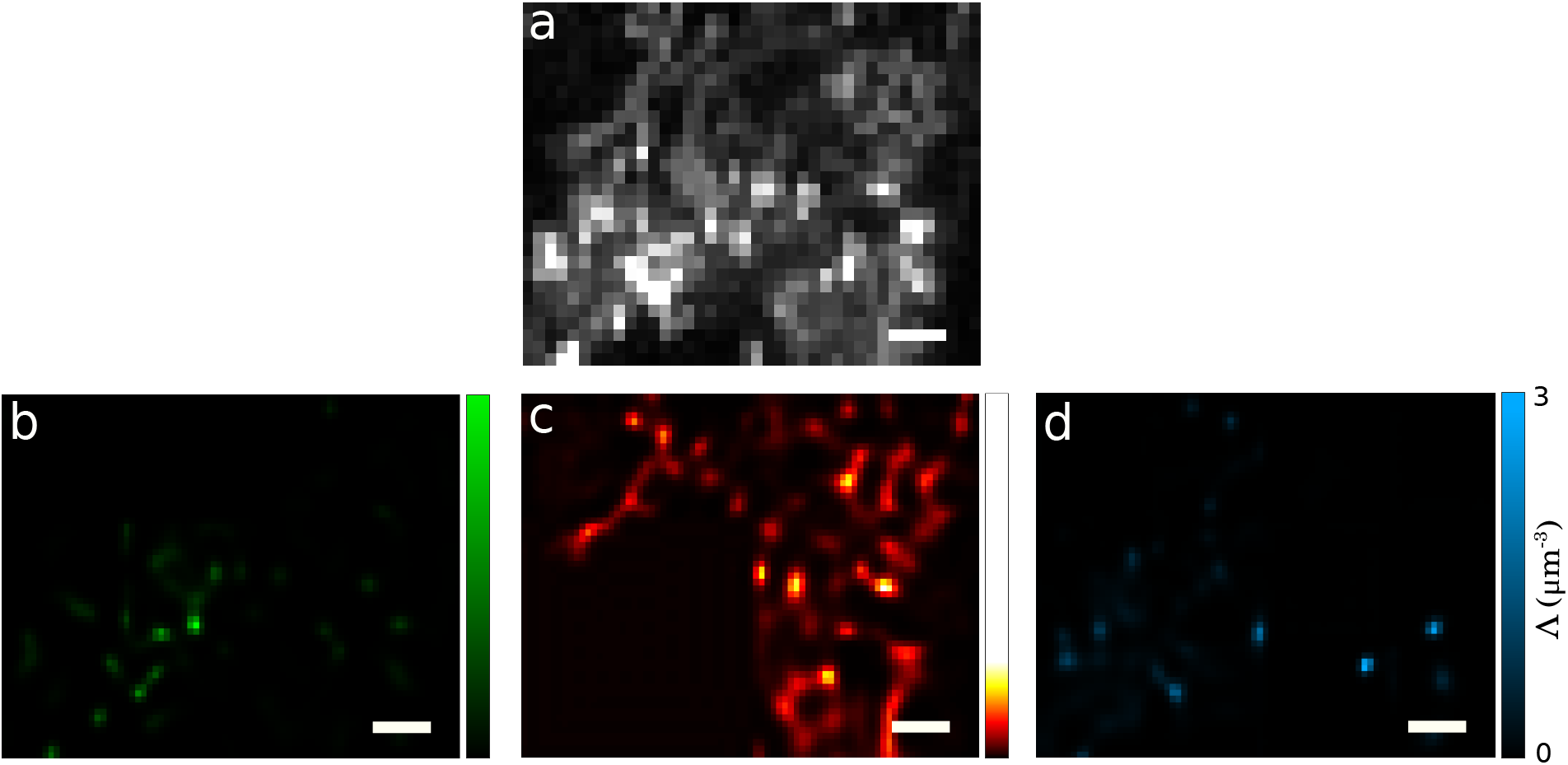
*In vivo* data set containing three lifetime components. (a) Data acquired by scanning the sample over area of 30×40 pixel. The sample is simultaneously labeled with three fluorophore species of pHrodo with lifetime of 0.8 ns staining lysosomes, TMRM with lifetime of 2.8 ns staining mitochondria, and Lyso-Red with lifetime of 4.5 ns staining endosomes. This resulted in three lifetime maps interpolated below pixel size (1/2 pixel); (b) lifetime maps corresponding to lifetime of 0.8 ns; (c) lifetime map corresponding to lifetime of 2.8 ns; (d) lifetime map corresponding to lifetime of 4.5 ns. Scale bars are 2 *μm*.

**Figure 4:**
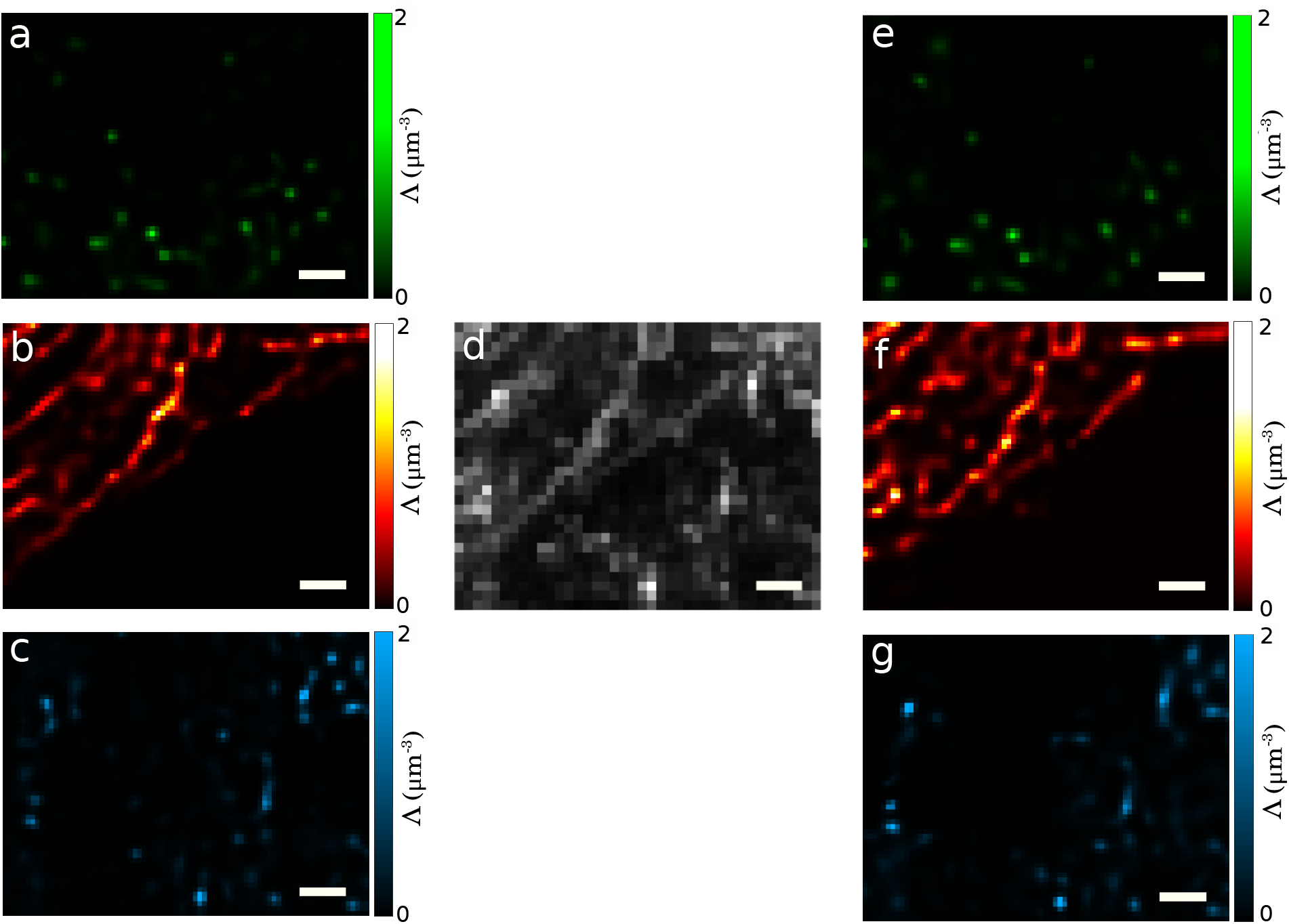
In order to test our method on realistic distributions of fluorophores, we consider *in vivo* FLIM data composed, as a test of our method, by mixing three single-species lifetime maps into one. That is, we first analyze three data sets each containing a single species to produce the “ground truth” maps seen in pnels a-c. More concretely: (a) is the “ground truth” lifetime map (green) for pHrodo with a lifetime of 0.8 ns; (b) is the “ground truth” lifetime map (red) for TMRM with a lifetime of 2.8 ns; and (c) is the “ground truth” lifetime map (blue) for Lyso-red with a lifetime of 4.5 ns. Now, we combine our three original data sets to produce (d). Next, we apply BNP-FLIM to learn the number of species and their maps that we show in panels e-g and can now compare, respectively, with panels a-c. Lifetime maps in panels a-c and e-g are reported with a pixel size equal to 1/2 the pixel size of panel d. Scale bars are 2 μm. The agreement between panels a-c and e-g is discussed in the text.

**Figure 5:**
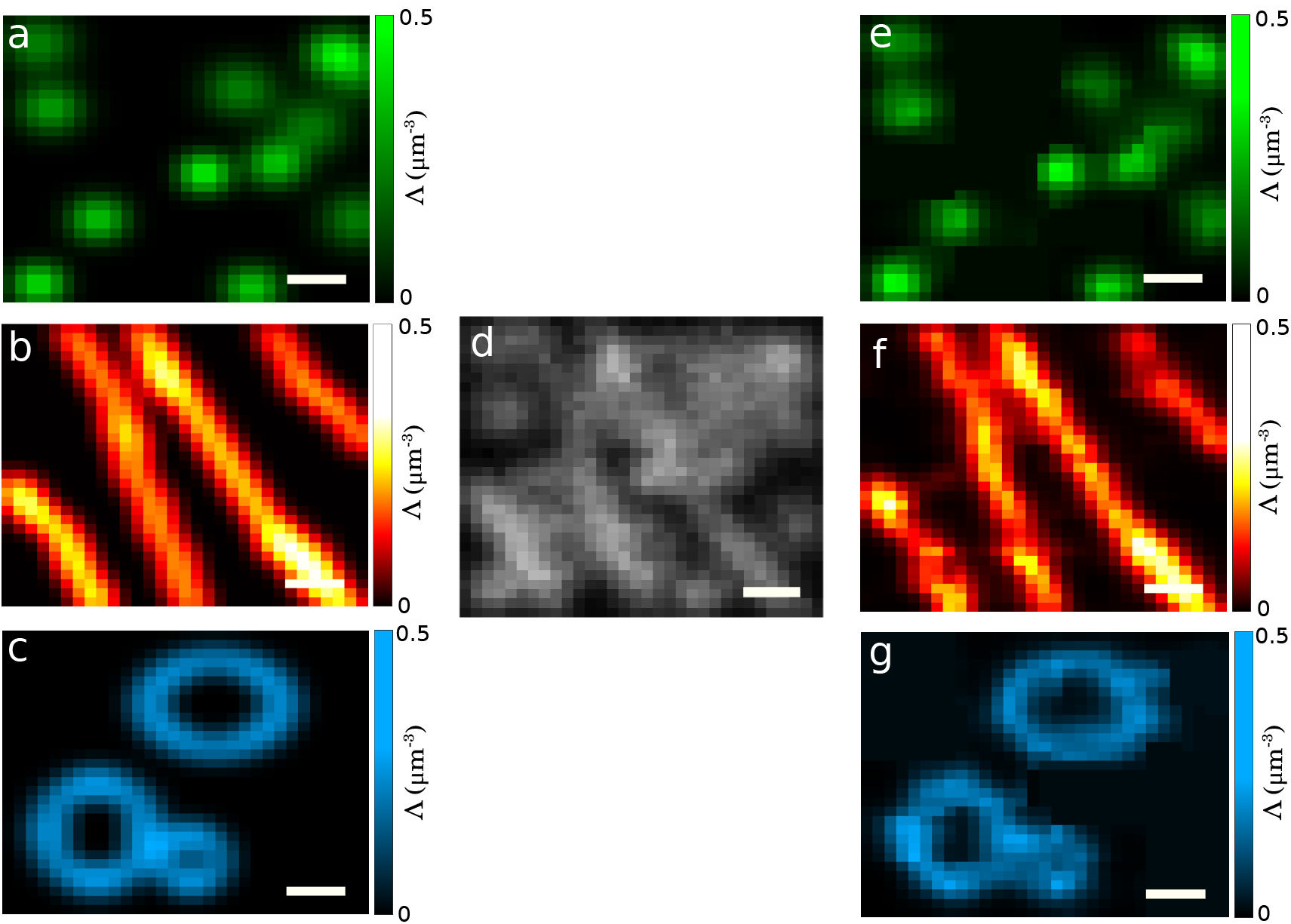
Simulated FLIM data generated from a mixture of three lifetime maps. (a) Ground truth map simulated with lifetime of 1 ns. (b) Ground truth map simulated with lifetime of 2.5 ns. (c) Ground truth map simulated with lifetime of 4.5 ns. (d) Data generated using a mixture of lifetime maps in panels a-c. This data was processed using the BNP-FLIM algorithm resulting again in three non-zero binary weights and corresponding lifetime maps with lifetimes of (e) 1 ns, (f) 2.5 ns, (g) 4.5 ns. Scale bars are 2 *μm*.

Before describing our results, we illustrate our procedure for simulating data. To be more precise, we explain the data simulation procedure for a single pixel (*i*th pixel) in order to later simulate all multi-pixel FLIM data used across this study. To do so, we assume a set of 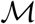 lifetime maps corresponding to 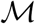 lifetime components (typically 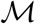 is much smaller than the truncated *M* we use in the Methods section to approximate the beta-Bernoulli process prior). Next, we use lifetime maps to simulate data in the *i*th pixel by: 1) sampling 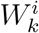 to determine whether the *k*th pulse results in a photon or not (*i.e*., is empty or nonempty pulse) from a Bernoulli distribution similar to eq. 2; 2) sampling the fluorophore species, 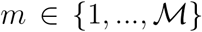, giving rise to the detected photons, for non-empty pulses, from a categorical distribution, *i.e*., a discrete distribution with more than two options, with probabilities 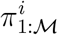 given by eqs. 4–5; 3) sampling the period spent in the excited state by the fluorophore, 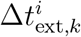, from an exponential distribution with lifetime *τ_m_*; 4) sampling the IRF time, 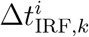, from a Gaussian distribution and adding it to the time spent in the excited state by the fluorophore; finally 5) deal with cases where the sum of both those times generated in steps 3-4 exceeds the interpulse time *T*.

To elaborate briefly on step 5, since the photon arrival times are recorded with respect to the immediately preceding pulse (sometimes termed the “microtime” in FLIM), they are always smaller than the interpulse window. As such, to allow for the possibility that some arrival times may exceed the interpulse window (especially when the excited state lifetime is on par with the interpulse window), we need to introduce a third term subtracting integer interpulse windows from the generated arrival times

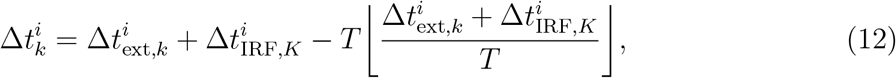

where the bracket returns the integer part of its content. We note, as a sanity check, that if an arrival time is smaller than the interpulse window the third term is zero. Moreover, the parameters used in the simulations are inspired by values from the experimental data used later. That is, we sample IRF times from a Gaussian distribution with mean and standard deviation of 12.20 ns and 0.8 ns. Furthermore, we use interpulse time and pixel size of *T* = 12.8 ns and 0.39 *μm* and assume a Gaussian PSF (see eq. 5) with *σ_xy_* = 0.54 *μm* and *σ_z_* = 1.56 μm. Other parameter values used in the analyses are provided in Supplementary Note 3.

### Robustness with respect to photon counts and lifetime resolution

Here, we start by considering smoothly varying lifetime maps (before turning to more complex maps in the next section). We use these maps to benchmark our BNP-FLIM algorithm against a range of lifetimes, lifetime differences, and photon counts. To do so, we use three synthetic maps corresponding to three different lifetimes over an area of 5 × 20 pixels. We use the combination of these maps to generate data sets with 10, 20 and 50 photons per pixel shown in the first, second and third columns in Fig. 2, respectively. Row a in Fig. 2 shows the resulting lifetime histograms where two of the lifetimes differ by only 0.3 ns representing a challenging case with sub-nanosecond lifetime differences. Here, the BNP-FLIM algorithm correctly assigns non-zero binary weights to three lifetime maps and learns the corresponding lifetimes accomplishing sub-nanosecond lifetime resolution even with 10 photons per pixel and 1000 photons in total. We also note that by increasing the photon counts to 20 and 50 photons per pixel the histograms’ widths tend to decrease signifying smaller uncertainties. Next, as we move to larger differences in lifetime in rows b-c, we expect to observe sharper histograms due to less uncertainty. However, the histograms do not exhibit such behavior due to posterior broadening for larger lifetimes. This is because fluorophore species with larger lifetimes are more likely to stay excited longer than an interpulse period leading to more photons detected one or more pulses following the one inducing fluorophore excitation. Depsite this, we still see histogram sharpening with more photons per pixel from left to right for each row. Further, cross-sections of lifetime maps corresponding to these lifetimes are represented in Fig. S1. Here, although we are able to achieve sub-nanosecond lifetime resolution by using photons from across the entire region, the spatial lifetime maps are learned using only photons from specific areas, *i.e*., pixels, which can result in poorer uncertainties in the estimated maps, particularly for the case with 10 photons per pixel, compared to the inferred lifetimes.

### Heterogeneous simulated and *in vivo* lifetime maps

In this section, we challenge our BNP-FLIM algorithm using lifetime maps with more complex features. To do so, we use data including: *in vivo* data acquired by labeling three different subcellular structures using different fluorophore species in Fig. 3; a mixed image composed of single-species *in vivo* data in Fig. 4 and Fig. S3-S6; and synthetic data generated from heterogeneous lifetime maps mimicking ones recovered *in vivo* (see Fig. 5).

Here, we start by assessing the performance of the BNP-FLIM algorithm using data acquired by scanning an area of 30× 40 pixels of an *in vivo* sample stained by three fluorophore species: pHrodo with a lifetime of 0.8 ns which binds to lysosomes; TMRM with a lifetime of 2.8 ns which binds to mitochondria; Lyso-red with a lifetime of 4.5 which binds to endosomes (see Fig. 3a). The BNP-FLIM algorithm correctly learns the number of lifetimes present in the data with the corresponding lifetimes shown in Fig. S2. For the lifetimes, the average difference of the ground truths and the histogram peaks is approximately 0.2 ns (where the ground truth in the SI is taken to originate from a very large number of photons using the phasor method^33^). Moreover, Fig. 3 depicts lifetime maps interpolated below pixel size (1/2 pixel) corresponding to pHrodo, TMRM and Lyso-Red shown in green, red and blue, respectively. However, in this case, we cannot benchmark BNP-FLIM using lifetime maps due to the lack of ground truth maps. For this reason, we use a hybrid data set consisting of three single species *in vivo* data and synthetic data described below.

We begin by first describing each single-species lifetime map within the hybrid *in vivo* data set. These data sets were acquired by scanning 30×40 pixel regions of three different samples (see Fig. S3a-c). Each sample contained one of the fluorophore species: pHrodo, TMRM and Lyso-red. Fig. 4a-c, respectively, represents the lifetime maps learned from the raw data shown in Fig. S3 reporting lifetime maps at 1/2 the pixel for pHrodo (green), TMRM (red) and Lyso-Red (blue). These lifetime maps are obtained by analyzing each individual data set using the BNP-FLIM algorithm (also see Fig. S3d-f). Moreover, Fig. S4 shows the resulting lifetime histograms where the histogram peaks differ from the ground truth by less than 0.07 ns on average (where ground truth lifetimes were again taken to be values found using phasor method).

We now evaluate the performance of our algorithm by combining all three data sets. The ground truth is now taken as the lifetime maps shown in Fig. 4a-c that were obtained by analyzing the single-species data sets.

Our BNP-FLIM algorithm correctly identified three species (as the maximum a posteriori estimate) and their coinciding lifetime maps are shown in Fig. 4e-g with the learned maps interpolated below pixel size (1/2 pixel) for pHrodo (green), TMRM (red) and Lyso-Red (blue). To quantitatively compare the resulting maps in Fig. 4a-c and the ground truth maps (Fig. 4e-g), we calculated their absolute relative differences given by |ΔΛ|/max(Λ_true_), where Λ_true_ denotes the ground truth map. The error maps are shown in Fig. S5 with the mean errors of ≈ 3%, ≈ 8% and ≈ 6% for pHrodo, TMRM and Lyso-Red, respectively. Moreover, the resulting histograms of the number of non-zero binary weights and the corresponding lifetimes are shown in Fig. S6. For lifetimes, the average difference of ground truth values and histogram peaks is approximately 0.12 ns.

After assessing the performance of our BNP-FLIM algorithm using *in vivo* data, we continue to benchmark our algorithm using synthetic FLIM data containing three complex lifetime maps (see Fig. 5a-c) similar to those in experimental data with lifetimes of 1 ns, 2.5 ns and 4.5 ns, generated over an area of 32×32 pixels (see Fig. 5d). We used our BNP-FLIM algorithm to analyze this data. The resulting lifetime maps are shown in Fig. 5e-g with average absolute relative differences of ≈ 5%, ≈ 8% and ≈ 5%, respectively. Moreover, the histograms of the number of non-zero binary weights warranted by the data and the corresponding lifetimes are given in Fig. S7. Here, the histogram in the number of lifetime components have peaks at the true value of 3 and peaks of the lifetime histograms differ from the ground truths by less than 0.08 ns on average.

## Discussion

Fluorescence lifetime imaging techniques allow us to probe life within complex sub-cellular environments at the nanoscale. In particular, these techniques have been employed to detect changes in cellular metabolism due to cancer metastasis^29,79,80^ whose ensuing metabolic shift is detected by monitoring varying levels of free and bound NADH within breast tissue cells^29^. However, quantitative interpretation of FLIM data remain limited by fundamentally unknown numbers of lifetime components to which to fit the data, as well as low lifetime and spatial resolutions, and high photon counts required to determine lifetimes separated by small differences. Addressing these issues requires a framework leveraging all spatial correlations across all pixels known physics of the problem by incorporating it directly into the likelihood. This includes Poisson photon emission statistics as well as all existing noise sources, *e.g*., IRF, to simultaneously learn the number of lifetime components and associated maps interpolated below data pixel size leveraging all spatial correlations across pixels.

Due to our choice of Monte Carlo samplers in BNP-FLIM, the computational cost scales linearly with the number of pixels and photon counts. Moreover, the computational cost also depends on the type of data analyzed. For instance, more challenging data sets, such as data sets containing lifetime maps with more complex features or multiple lifetime maps (where background could be considered yet another species), complicate the posterior’s shape. This necessarily requires us to draw more samples from the posterior and therefore introduces higher computational cost. For example, it took two days to analyze the single species *in vivo* data sets shown in Fig. S3, while it took about a week to analyze the *in vivo* data set containing three lifetime maps shown in Fig. 4d on a regular desktop machine.

Although we assumed a Gaussian IRF for simplicity while retaining sufficient accuracy, the BNP-FLIM algorithm is also capable of accommodating any type of IRF by simply modifying eq. 3 at no additional computational cost. Moreover, here, we assumed that parameters of the IRF distribution, *e.g*., offset and variance, can be pre-calibrated. However, our inverse strategy can be generalized to learn these parameters along the rest of unknowns by adding appropriate priors on these parameters. What is more, for species present at low concentrations, the probability of no excitation, appearing in eq. 5, can be approximated by the first two terms of their Taylor expansion, leading to simpler expressions and reduced computational complexity.

## References

(1) Suhling, K.; Hirvonen, L. M.; Levitt, J. A.; Chung, P.-H.; Tregidgo, C.; Le Marois, A.; Rusakov, D. A.; Zheng, K.; Ameer-Beg, S.; Poland, S., et al. Fluorescence lifetime imaging (FLIM): Basic concepts and some recent developments. Medical photonics 2015, 27, 3–40.

(2) Datta, R.; Heaster, T. M.; Sharick, J. T.; Gillette, A. A.; Skala, M. C. Fluorescence lifetime imaging microscopy: fundamentals and advances in instrumentation, analysis, and applications. Journal of biomedical optics 2020, 25, 071203.

(3) Garini, Y.; Young, I. T.; McNamara, G. Spectral imaging: principles and applications. Cytometry part a: the journal of the international society for analytical cytology 2006, 69, 735–747.

(4) Huang, B.; Bates, M.; Zhuang, X. Super-resolution fluorescence microscopy. Annual review of biochemistry 2009, 78, 993–1016.

(5) Lelek, M.; Gyparaki, M. T.; Beliu, G.; Schueder, F.; Griffié, J.; Manley, S.; Jungmann, R.; Sauer, M.; Lakadamyali, M.; Zimmer, C. Single-molecule localization microscopy. Nature reviews methods primers 2021, 1, 1–27.

(6) Fazel, M.; Wester, M. J. Analysis of super-resolution single molecule localization microscopy data: A tutorial. AIP advances 2022, 12, 010701.

(7) Lidke, D.; Nagy, P.; Barisas, B.; Heintzmann, R.; Post, J. N.; Lidke, K.; Clayton, A.; Arndt-Jovin, D.; Jovin, T. Imaging molecular interactions in cells by dynamic and static fluorescence anisotropy (rFLIM and emFRET). Biochemical society transactions 2003, 31, 1020–1027.

(8) Tregidgo, C. L.; Levitt, J. A.; Suhling, K. Effect of refractive index on the fluorescence lifetime of green fluorescent protein. Journal of biomedical optics 2008, 13, 031218.

(9) Coban, O.; Zanetti-Dominguez, L. C.; Matthews, D. R.; Rolfe, D. J.; Weitsman, G.; Barber, P. R.; Barbeau, J.; Devauges, V.; Kampmeier, F.; Winn, M., et al. Effect of phosphorylation on EGFR dimer stability probed by single-molecule dynamics and FRET/FLIM. Biophysical journal 2015, 108, 1013–1026.

(10) Aubret, A.; Pillonnet, A.; Houel, J.; Dujardin, C.; Kulzer, F. CdSe/ZnS quantum dots as sensors for the local refractive index. Nanoscale 2016, 8, 2317–2325.

(11) Pliss, A.; Prasad, P. N. High resolution mapping of subcellular refractive index by Fluorescence Lifetime Imaging: a next frontier in quantitative cell science? Methods and applications in fluorescence 2020, 8, 032001.

(12) Joosen, L.; Hink, M.; Gadella Jr, T.; Goedhart, J. Effect of fixation procedures on the fluorescence lifetimes of Aequorea victoria derived fluorescent proteins. Journal of microscopy 2014, 256, 166–176.

(13) Inada, N.; Fukuda, N.; Hayashi, T.; Uchiyama, S. Temperature imaging using a cationic linear fluorescent polymeric thermometer and fluorescence lifetime imaging microscopy. Nature protocols 2019, 14, 1293–1321.

(14) Kalytchuk, S.; Polakova, K.; Wang, Y.; Froning, J. P.; Cepe, K.; Rogach, A. L.; Zboril, R. Carbon dot nanothermometry: intracellular photoluminescence lifetime thermal sensing. ACS nano 2017, 11, 1432–1442.

(15) Okabe, K.; Inada, N.; Gota, C.; Harada, Y.; Funatsu, T.; Uchiyama, S. Intracellular temperature mapping with a fluorescent polymeric thermometer and fluorescence lifetime imaging microscopy. Nature communications 2012, 3, 1–9.

(16) Zhang, H.; Jiang, J.; Gao, P.; Yang, T.; Zhang, K. Y.; Chen, Z.; Liu, S.; Huang, W.; Zhao, Q. Dual-emissive phosphorescent polymer probe for accurate temperature sensing in living cells and zebrafish using ratiometric and phosphorescence lifetime imaging microscopy. ACS applied materials & interfaces 2018, 10, 17542–17550.

(17) Yin, J.; Peng, M.; Lin, W. Visualization of mitochondrial viscosity in inflammation, fatty liver, and cancer living mice by a robust fluorescent probe. Analytical chemistry 2019, 91, 8415–8421.

(18) Hao, L.; Li, Z.-W.; Zhang, D.-Y.; He, L.; Liu, W.; Yang, J.; Tan, C.-P.; Ji, L.-N.; Mao, Z.-W. Monitoring mitochondrial viscosity with anticancer phosphorescent Ir (III) complexes via two-photon lifetime imaging. Chemical science 2019, 10, 1285–1293.

(19) Liu, T.; Campbell, B. T.; Burns, S. P.; Sullivan, J. P., et al. Temperature-and pressure-sensitive luminescent paints in aerodynamics. Applied mechanics reviews 1997, 50, 227–246.

(20) Gregory, J.; Asai, K.; Kameda, M.; Liu, T.; Sullivan, J. A review of pressure-sensitive paint for high-speed and unsteady aerodynamics. Proceedings of the institution of mechanical engineers, part G: journal of aerospace engineering 2008, 222, 249–290.

(21) Lin, H.-J.; Herman, P.; Lakowicz, J. R. Fluorescence lifetime-resolved pH imaging of living cells. Cytometry part A: the journal of the international society for analytical cytology 2003, 52, 77–89.

(22) Bizzarri, R.; Serresi, M.; Luin, S.; Beltram, F. Green fluorescent protein based pH indicators for in vivo use: a review. Analytical and bioanalytical chemistry 2009, 393, 1107–1122.

(23) Zhu, X.-H.; Lu, M.; Lee, B.-Y.; Ugurbil, K.; Chen, W. In vivo NAD assay reveals the intracellular NAD contents and redox state in healthy human brain and their age dependences. Proceedings of the national academy of sciences 2015, 112, 2876–2881.

(24) Bird, D. K.; Yan, L.; Vrotsos, K. M.; Eliceiri, K. W.; Vaughan, E. M.; Keely, P. J.; White, J. G.; Ramanujam, N. Metabolic mapping of MCF10A human breast cells via multiphoton fluorescence lifetime imaging of the coenzyme NADH. Cancer research 2005, 65, 8766–8773.

(25) Ma, N.; Digman, M. A.; Malacrida, L.; Gratton, E. Measurements of absolute concentrations of NADH in cells using the phasor FLIM method. Biomedical optics express 2016, 7, 2441–2452.

(26) Frei, M. S.; Tarnawski, M.; Roberti, M. J.; Koch, B.; Hiblot, J.; Johnsson, K. Engineered HaloTag variants for fluorescence lifetime multiplexing. Nature methods 2022, 19, 65–70.

(27) Blacker, T. S.; Mann, Z. F.; Gale, J. E.; Ziegler, M.; Bain, A. J.; Szabadkai, G.; Duchen, M. R. Separating NADH and NADPH fluorescence in live cells and tissues using FLIM. Nature communications 2014, 5, 1–9.

(28) Homulle, H.; Powolny, F.; Stegehuis, P.; Dijkstra, J.; Li, D.-U.; Homicsko, K.; Rimoldi, D.; Muehlethaler, K.; Prior, J.; Sinisi, R., et al. Compact solid-state CMOS single-photon detector array for in vivo NIR fluorescence lifetime oncology measurements. Biomedical optics express 2016, 7, 1797–1814.

(29) Davis, R. T.; Blake, K.; Ma, D.; Gabra, M. B. I.; Hernandez, G. A.; Phung, A. T.; Yang, Y.; Maurer, D.; Lefebvre, A. E.; Alshetaiwi, H., et al. Transcriptional diversity and bioenergetic shift in human breast cancer metastasis revealed by single-cell RNA sequencing. Nature cell biology 2020, 22, 310–320.

(30) Skala, M. C.; Riching, K. M.; Bird, D. K.; Gendron-Fitzpatrick, A.; Eickhoff, J.; Eliceiri, K. W.; Keely, P. J.; Ramanujam, N. In vivo multiphoton fluorescence lifetime imaging of protein-bound and free nicotinamide adenine dinucleotide in normal and precancerous epithelia. Journal of biomedical optics 2007, 12, 024014.

(31) Kumar, S.; Alibhai, D.; Margineanu, A.; Laine, R.; Kennedy, G.; McGinty, J.; Warren, S.; Kelly, D.; Alexandrov, Y.; Munro, I., et al. FLIM FRET Technology for Drug Discovery: Automated Multiwell-Plate High-Content Analysis, Multiplexed Readouts and Application in Situ. ChemPhysChem 2011, 12, 609–626.

(32) Conway, J. R.; Carragher, N. O.; Timpson, P. Developments in preclinical cancer imaging: innovating the discovery of therapeutics. Nature reviews cancer 2014, 14, 314–328.

(33) Digman, M. A.; Caiolfa, V. R.; Zamai, M.; Gratton, E. The phasor approach to fluorescence lifetime imaging analysis. Biophysical journal 2008, 94, L14–L16.

(34) Ranjit, S.; Malacrida, L.; Jameson, D. M.; Gratton, E. Fit-free analysis of fluorescence lifetime imaging data using the phasor approach. Nature protocols 2018, 13, 1979–2004.

(35) Vallmitjana, A.; Torrado, B.; Dvornikov, A.; Ranjit, S.; Gratton, E. Blind resolution of lifetime components in individual pixels of fluorescence lifetime images using the phasor approach. The journal of physical chemistry B 2020, 124, 10126–10137.

(36) Scipioni, L.; Rossetta, A.; Tedeschi, G.; Gratton, E. Phasor S-FLIM: a new paradigm for fast and robust spectral fluorescence lifetime imaging. Nature methods 2021, 18, 542–550.

(37) Wu, G.; Nowotny, T.; Zhang, Y.; Yu, H.-Q.; Li, D. D.-U. Artificial neural network approaches for fluorescence lifetime imaging techniques. Optics letters 2016, 41, 2561–2564.

(38) Smith, J. T.; Yao, R.; Sinsuebphon, N.; Rudkouskaya, A.; Un, N.; Mazurkiewicz, J.; Barroso, M.; Yan, P.; Intes, X. Fast fit-free analysis of fluorescence lifetime imaging via deep learning. Proceedings of the national academy of sciences 2019, 116, 24019–24030.

(39) Yao, R.; Ochoa, M.; Yan, P.; Intes, X. Net-FLICS: fast quantitative wide-field fluorescence lifetime imaging with compressed sensing–a deep learning approach. Light: science & applications 2019, 8, 1–7.

(40) Verveer, P. J.; Squire, A.; Bastiaens, P. I. Global analysis of fluorescence lifetime imaging microscopy data. Biophysical journal 2000, 78, 2127–2137.

(41) Straume, M.; Frasier-Cadoret, S. G.; Johnson, M. L. Topics in fluorescence spectroscopy; Springer, 2002; pp 177–240.

(42) Pelet, S.; Previte, M.; Laiho, L.; So, P. A fast global fitting algorithm for fluorescence lifetime imaging microscopy based on image segmentation. Biophysical journal 2004, 87, 2807–2817.

(43) Bajzer, Ž.; Therneau, T. M.; Sharp, J. C.; Prendergast, F. G. Maximum likelihood method for the analysis of time-resolved fluorescence decay curves. European biophysics journal 1991, 20, 247–262.

(44) Maus, M.; Cotlet, M.; Hofkens, J.; Gensch, T.; De Schryver, F. C.; Schaffer, J.; Seidel, C. An experimental comparison of the maximum likelihood estimation and nonlinear least-squares fluorescence lifetime analysis of single molecules. Analytical chemistry 2001, 73, 2078–2086.

(45) Thiele, J. C.; Nevskyi, O.; Helmerich, D. A.; Sauer, M.; Enderlein, J. Advanced data analysis for Fluorescence-Lifetime Single-Molecule Localization Microscopy. Frontiers in bioinformatics 2021, 56.

(46) Barber, P. R.; Ameer-Beg, S. M.; Pathmananthan, S.; Rowley, M.; Coolen, A. A Bayesian method for single molecule, fluorescence burst analysis. Biomedical optics express 2010, 1, 1148–1158.

(47) Rowley, M. I.; Barber, P. R.; Coolen, A. C.; Vojnovic, B. Bayesian analysis of fluorescence lifetime imaging data. Multiphoton Microscopy in the Biomedical Sciences XI. 2011; p 790325.

(48) Rowley, M. I.; Coolen, A. C.; Vojnovic, B.; Barber, P. R. Robust Bayesian fluorescence lifetime estimation, decay model selection and instrument response determination for low-intensity FLIM imaging. PLoS one 2016, 11, e0158404.

(49) Kaye, B.; Foster, P. J.; Yoo, T. Y.; Needleman, D. J. Developing and testing a bayesian analysis of fluorescence lifetime measurements. PLoS one 2017, 12, e0169337.

(50) Wang, S.; Chacko, J. V.; Sagar, A. K.; Eliceiri, K. W.; Yuan, M. Nonparametric empirical Bayesian framework for fluorescence-lifetime imaging microscopy. Biomedical optics express 2019, 10, 5497–5517.

(51) Santra, K.; Smith, E. A.; Song, X.; Petrich, J. W. A Bayesian approach for extracting fluorescence lifetimes from sparse data sets and its significance for imaging experiments. Photochemistry and photobiology 2019, 95, 773–779.

(52) Fazel, M.; Vallmitjana, A.; Scipioni, L.; Gratton, E.; Digman, M. A.; Presse, S. Fluorescence Lifetime: Beating the IRF and interpulse window. bioRxiv 2022,

(53) Ware, W. R.; Doemeny, L. J.; Nemzek, T. L. Deconvolution of fluorescence and phosphorescence decay curves. Least-squares method. The journal of physical chemistry 1973, 77, 2038–2048.

(54) Jo, J. A.; Fang, Q.; Papaioannou, T.; Marcu, L. Laguerre nonparametric deconvolution technique of time-resolved fluorescence data: application to the prediction of concentrations in a mixture of biochemical components. Optical biopsy V. 2004; pp 8–16.

(55) Campos-Delgado, D. U.; Navarro, O. G.; Arce-Santana, E.; Walsh, A. J.; Skala, M. C.; Jo, J. A. Deconvolution of fluorescence lifetime imaging microscopy by a library of exponentials. Optics express 2015, 23, 23748–23767.

(56) Tavakoli, M.; Jazani, S.; Sgouralis, I.; Heo, W.; Ishii, K.; Tahara, T.; Pressé, S. Direct Photon-by-Photon Analysis of Time-Resolved Pulsed Excitation Data using Bayesian Nonparametrics. Cell reports physical science 2020, 1, 100234.

(57) Le Marois, A.; Labouesse, S.; Suhling, K.; Heintzmann, R. Noise-Corrected Principal Component Analysis of fluorescence lifetime imaging data. Journal of biophotonics 2017, 10, 1124–1133.

(58) Fazel, M.; Jazani, S.; Scipioni, L.; Vallmitjana, A.; Gratton, E.; Digman, M. A.; Pressé, S. High resolution fluorescence lifetime maps from minimal photon counts. ACS photonics 2022, 9, 1015–1025.

(59) Stringari, C.; Cinquin, A.; Cinquin, O.; Digman, M. A.; Donovan, P. J.; Gratton, E. Phasor approach to fluorescence lifetime microscopy distinguishes different metabolic states of germ cells in a live tissue. Proceedings of the national academy of sciences 2011, 108, 13582–13587.

(60) Li, Y.; Natakorn, S.; Chen, Y.; Safar, M.; Cunningham, M.; Tian, J.; Li, D. D.-U. Investigations on average fluorescence lifetimes for visualizing multi-exponential decays. Frontiers in physics 2020, 8, 576862.

(61) Paisley, J.; Carin, L. Nonparametric factor analysis with beta process priors. Proceedings of the 26th annual international conference on machine learning. 2009; pp 777–784.

(62) Al Labadi, L.; Zarepour, M. On approximations of the beta process in latent feature models: Point processes approach. Sankhya A 2018, 80, 59–79.

(63) Jazani, S.; Sgouralis, I.; Shafraz, O. M.; Levitus, M.; Sivasankar, S.; Pressé, S. An alternative framework for fluorescence correlation spectroscopy. Nature communications 2019, 10, 1–10.

(64) Rasmussen, C. E. Gaussian processes in machine learning. Summer School on Machine Learning. 2003; pp 63–71.

(65) Quiñonero-Candela, J.; Rasmussen, C. E. A unifying view of sparse approximate Gaussian process regression. Journal of machine learning research 2005, 6, 1939–1959.

(66) Titsias, M. K.; Lawrence, N.; Rattray, M. Markov chain Monte Carlo algorithms for Gaussian processes. Inference and Estimation in Probabilistic Time-Series Models 2008, 9.

(67) Bryan IV, J. S.; Sgouralis, I.; Pressé, S. Inferring effective forces for Langevin dynamics using Gaussian processes. The journal of chemical physics 2020, 152, 124106.

(68) Sgouralis, I.; Pressé, S. An introduction to infinite HMMs for single-molecule data analysis. Biophysical journal 2017, 112, 2021–2029.

(69) Fazel, M.; Wester, M. J.; Mazloom-Farsibaf, H.; Meddens, M.; Eklund, A. S.; Schlichthaerle, T.; Schueder, F.; Jungmann, R.; Lidke, K. A. Bayesian multiple emitter fitting using reversible jump Markov Chain Monte Carlo. Scientific reports 2019, 9, 1–10.

(70) Fazel, M.; Wester, M. J.; Schodt, D. J.; Cruz, S. R.; Strauss, S.; Schueder, F.; Schlichthaerle, T.; Gillette, J. M.; Lidke, D. S.; Rieger, B., et al. High-precision estimation of emitter positions using Bayesian grouping of localizations. Nature Communications 2022, 13, 1–11.

(71) Tavakoli, M.; Jazani, S.; Sgouralis, I.; Shafraz, O. M.; Sivasankar, S.; Donaphon, B.; Levitus, M.; Pressé, S. Pitching single-focus confocal data analysis one photon at a time with Bayesian nonparametrics. Physical review X 2020, 10, 011021.

(72) Bryan IV, J. S.; Sgouralis, I.; Pressé, S. Diffraction-limited molecular cluster quantification with Bayesian nonparametrics. Nature computational science 2022, 2, 102–111.

(73) Saurabh, A.; Safar, M.; Sgouralis, I.; Fazel, M.; Pressé, S. Single photon smFRET. I. theory and conceptual basis. bioRxiv 2022,

(74) Saurabh, A.; Safar, M.; Fazel, M.; Sgouralis, I.; Pressé, S. Single photon smFRET. II. application to continuous illumination. bioRxiv 2022,

(75) Safar, M.; Saurabh, A.; Sarkar, B.; Fazel, M.; Ishii, K.; Tahara, T.; Sgouralis, I.; Presse, S. Single photon smFRET. III. application to pulsed illumination. bioRxiv 2022,

(76) Metropolis, N.; Rosenbluth, A. W.; Rosenbluth, M. N.; Teller, A. H.; Teller, E. Equation of state calculations by fast computing machines. The journal of chemical physics 1953, 21, 1087–1092.

(77) Hastings, W. K. Monte Carlo sampling methods using Markov chains and their applications. Biometrika 1970, 57, 97–109.

(78) Murray, I.; Adams, R.; MacKay, D. Elliptical slice sampling. Proceedings of the thirteenth international conference on artificial intelligence and statistics. 2010; pp 541–548.

(79) Ma, D.; Hernandez, G. A.; Lefebvre, A. E.; Alshetaiwi, H.; Blake, K.; Dave, K. R.; Rauf, M.; Williams, J. W.; Davis, R. T.; Evans, K. T., et al. Patient-derived xenograft culture-transplant system for investigation of human breast cancer metastasis. Communications biology 2021, 4, 1–15.

(80) Liu, L.; Zhang, S. X.; Liao, W.; Farhoodi, H. P.; Wong, C. W.; Chen, C. C.; Ségaliny, A. I.; Chacko, J. V.; Nguyen, L. P.; Lu, M., et al. Mechanoresponsive stem cells to target cancer metastases through biophysical cues. Science translational medicine 2017, 9, eaan2966.

