## Supplementary Information for "Building Fluorescence Lifetime Maps Photon-by-photon by Leveraging Spatial Correlations"

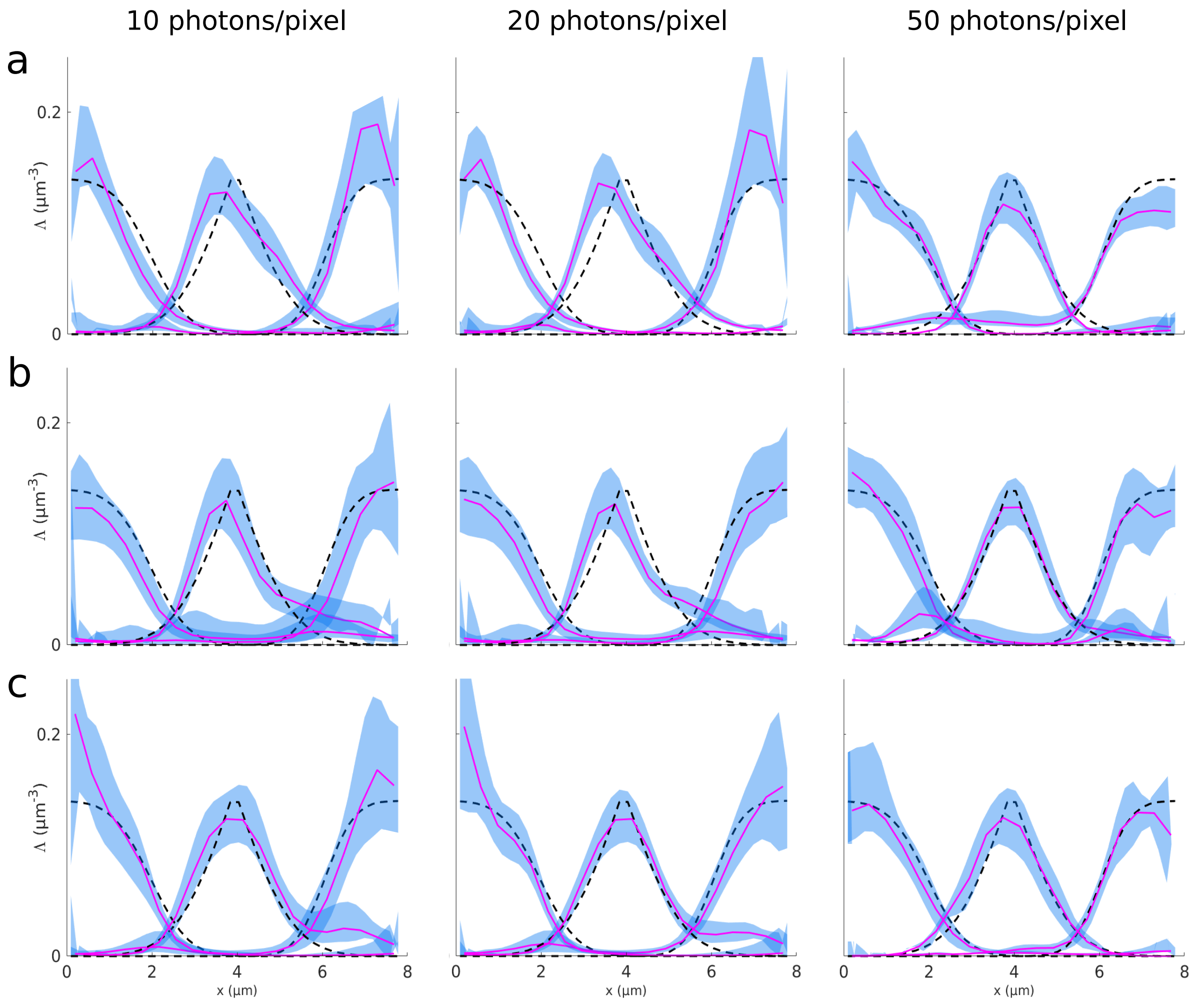


Fig. S1: BNP-FLIM robustness with respect to photon counts per pixel and lifetime differences. Three overlapping lifetime maps with different lifetimes were generated over a region of 5×20 pixels (pixel size of 0.39 $\mu m$) and analyzed by the BNP-FLIM algorithm. Columns from left to right, respectively, represent cross sections of the resulting lifetime maps with 10, 20 and 50 photons per pixel. All the data sets contain three lifetimes involving two lifetimes of 1 ns and 4.5 ns similar across all the generated data. The third lifetime varies: row (a) 1.3 ns; row (b) 1.8 ns; and row (c) 2.5 ns. The dashed curves, blue area and the magenta curves, respectively, show ground truths, 95% confidence interval of the sampled lifetime maps and the median of the sampled lifetime maps.

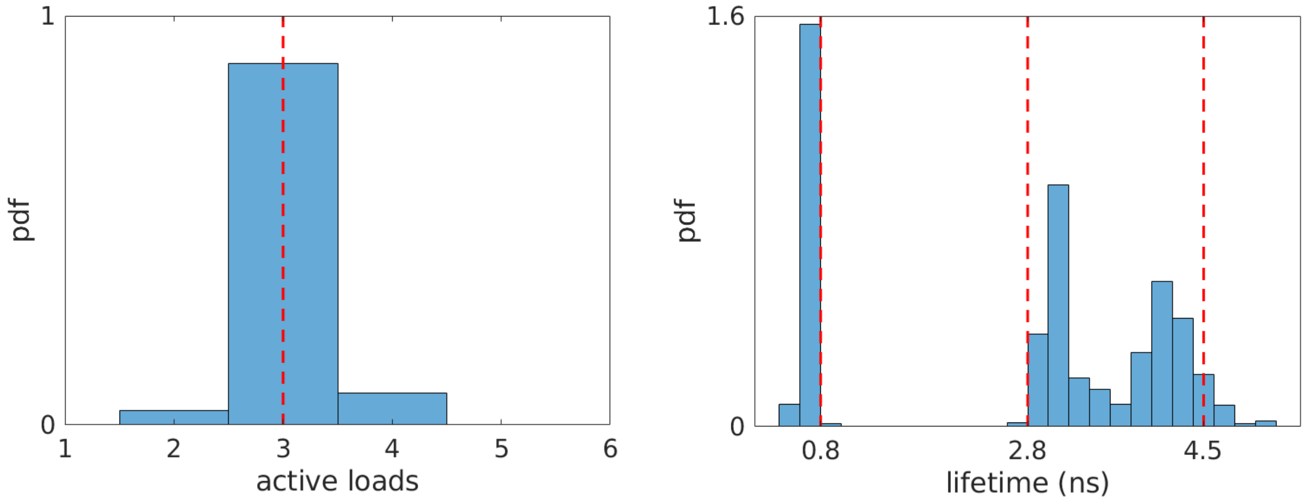


Fig. S2: Resulting histograms of the number of non-zero binary weights (loads) and the corresponding lifetimes from analyses of the simulated data shown in Fig. 3. Red dashed lines represent ground truths. We retain the same convention hereafter.

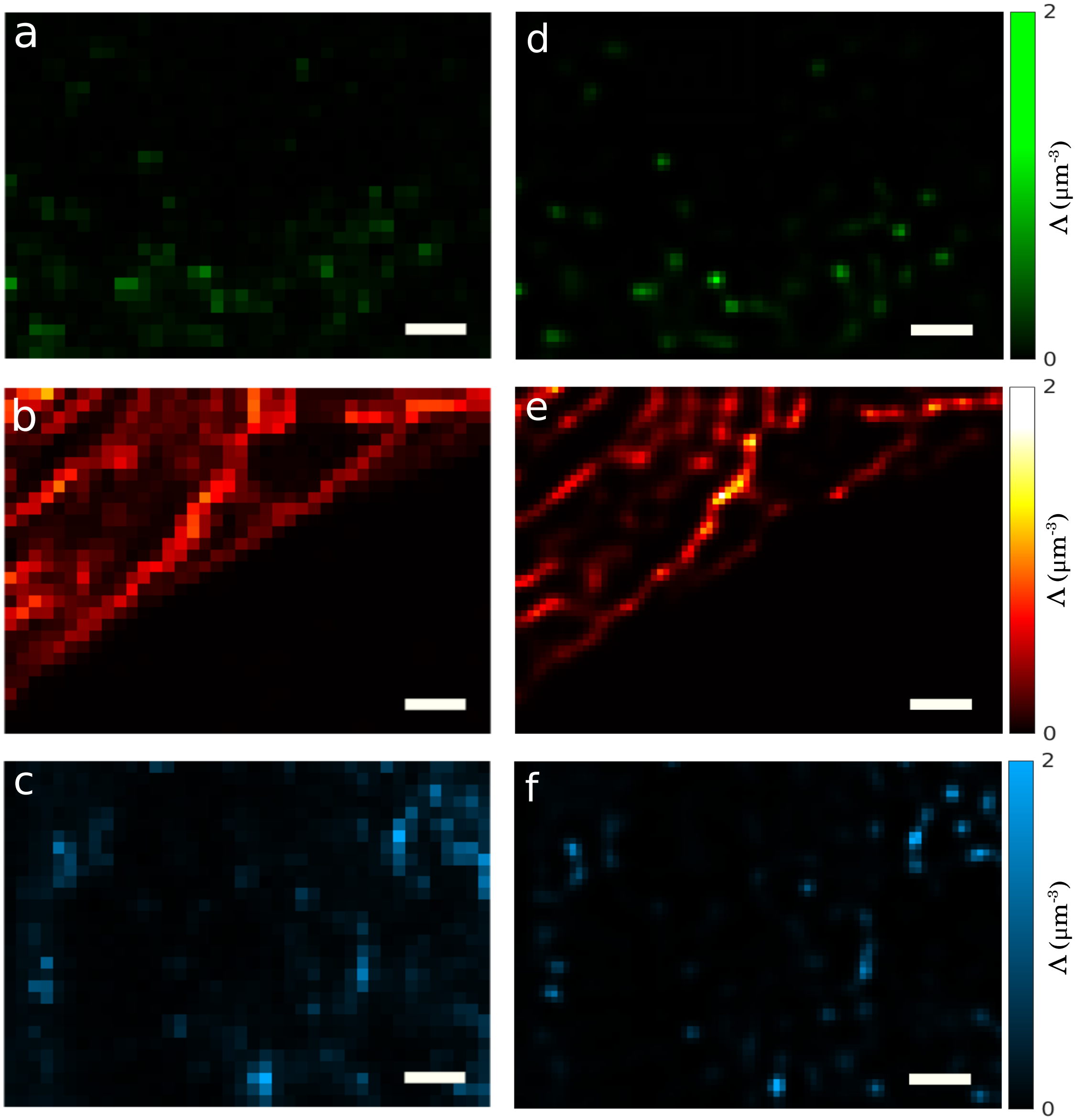


Fig. S3: *In vivo* data sets each containing a single lifetime component. (a) Data acquired using pHrodo fluorophores with lifetime of 0.8 ns labeling lysosomes. (b) Data acquired using TMRM fluorophores with lifetime of 2.8 ns labeling mitochondria. (c) Data acquired using Lyso-red fluorophores with lifetime of 4.5 ns labeling endosomes. These data sets (as well as other data sets acquired over large areas) were processed using the BNP-FLIM algorithm by splitting the entire region into multiple subregions and processing each subregion independently. The corresponding resulting lifetime maps are represented in panels d-f. Scale bars are 2 $\mu m$

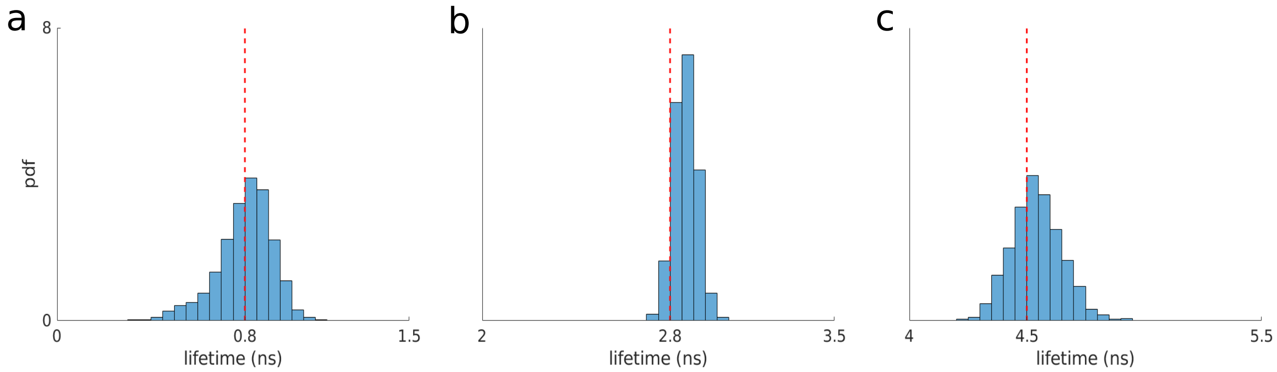


Fig. S4: Resulting lifetime histograms from analyses of data with single lifetimes in Fig. 3a-c.

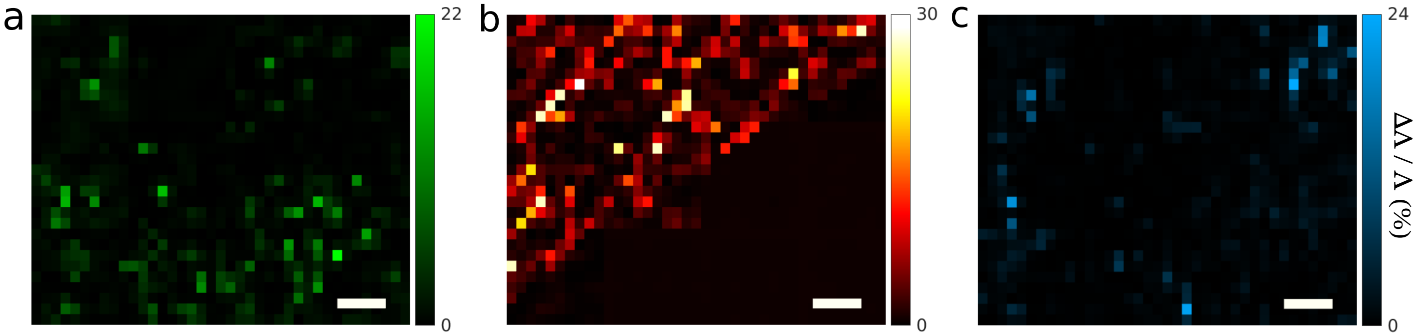


Fig. S5: Error maps given as the absolute relative differences of the discerned and ground truth lifetime maps shown in Fig. 4. (a) Error map corresponding to the lifetime map of pHrodo fluorophores with average error of ≈ 3%. (b) Error map corresponding to the lifetime map of TMRM fluorophores with average error of ≈ 8%. (c) Error map corresponding to the lifetime map of Lyso-red with average error of ≈ 6%.

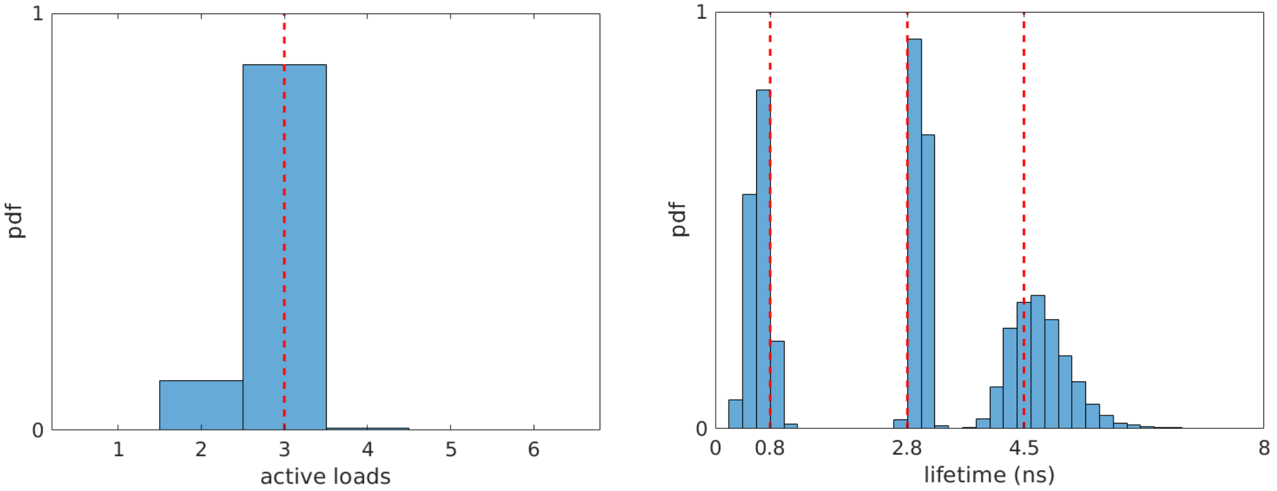


Fig. S6: Resulting histograms of the number of non-zero binary weights (loads) and the corresponding lifetimes from analyses of the *in vivo* data using mixture of three lifetime maps shown in Fig. 4.

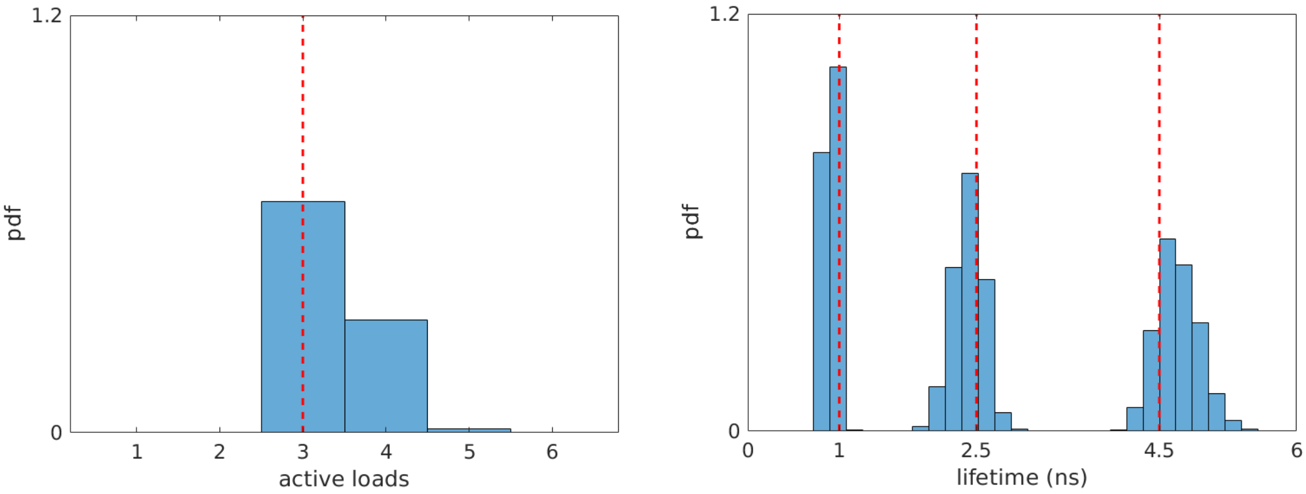


Fig. S7: Resulting histograms of the number of non-zero binary weights (loads) and the corresponding lifetimes from analyses of the simulated data using mixture of three lifetime maps shown in Fig. 5.

### Supplementary Note 1: Model Description

#### Supplementary Note 1.1: Likelihood

In this section, we present the mathematical model of Bayesian nonparametric FLIM (BNP-FLIM) in more detail. To do so, we start from the likelihood. The likelihood of BNP-FLIM is given by the product of the likelihoods of individual photon arrival times ($\Delta t_k^i$) and pulses ($W_k^i$) as follows

$$P\left(\overline{\overline{\Delta t}}, \overline{\overline{W}}|\vartheta\right) = \prod_i \prod_k P\left(\Delta t_k^i|\vartheta\right) P\left(W_k^i|\vartheta\right), \tag{S1}$$

where overlines denote the entire sets of arrival times and pulses. Moreover, $\vartheta = (\tau_{1:M}, \Lambda_{1:M}, \nu_{1:M}, b_{1:M})$ represents the set of unknown parameters we wish to infer: lifetimes, lifetime maps, means of Gaussian Process priors (GP), and binary weights. The likelihood for the $k$th pulse in the $i$th pixel leading to a photon observation or not (empty or non-empty pulse) is provided in the main text eq. 2 as

$$P\left(W_k^i|\vartheta\right) = \text{Bernoulli}\left(W_k^i; 1-\pi_0^i\right), \tag{S2}$$

where $\pi_0^i$ is the probability of no photon detection from the $i$th pixel. In what follows, we will derive the likelihood model for the arrival times.

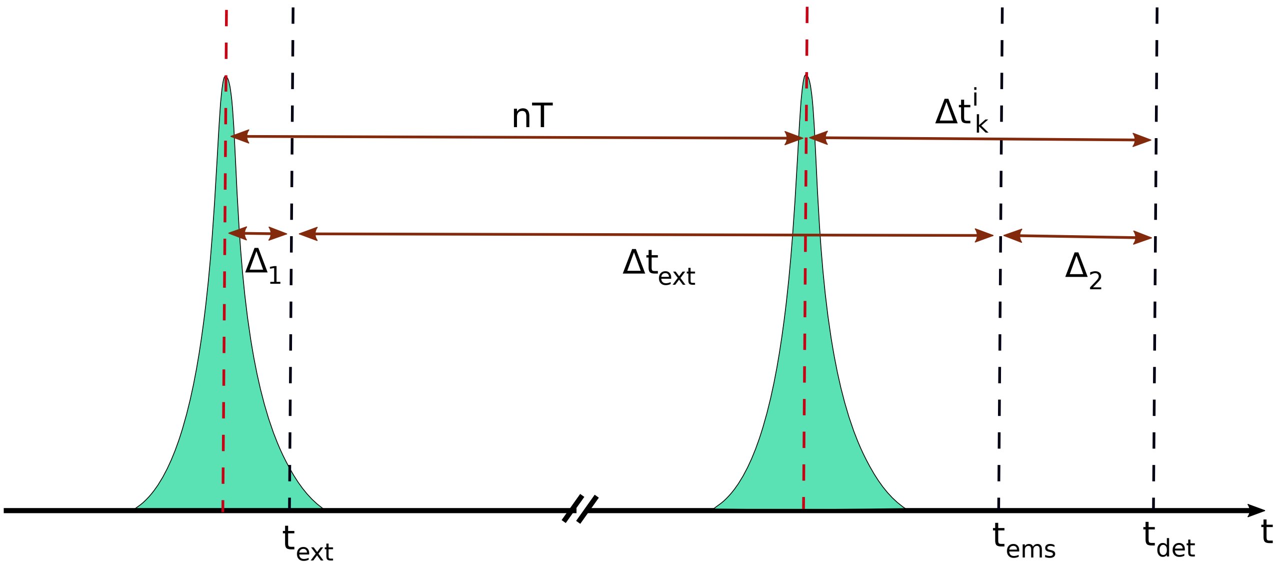


Fig. S8: Laser pulses and the arrival time likelihood. The green spikes and red dashed lines, respectively, represent the laser pulses and their centers. The black dashed lines show a fluorophore excitation time ($t_{\text{ext}}$), photon emission time ($t_{\text{ems}}$), and photon detection time ($t_{\text{det}}$). Here, $n, T, \Delta t_{\text{ext}}$ and $\Delta_k^i$ denote the number of pulses after which the photon emission takes place, the inter-pulse period, the time that the fluorophore spent in in the excited state, and the reported photon arrival time with respect to the immediate previous pulse. Moreover, $\Delta_1 + \Delta_2 = \Delta_{\text{IRF}}$ denote the instrument response function due to the delay in the detector in reporting the photon detection and the difference of the excitation time and the center of the pulse.

To derive the likelihood model for the arrival times, we start from the following expression obtained considering Fig. S8

$$\Delta t_k^i = \Delta t_{\text{ext}} + (\Delta_1 + \Delta_2) - nT = \Delta t_{\text{ext}} + \Delta t_{\text{IRF}} - nT. \tag{S3}$$

The above equation indicates that the observed photon arrival time (measurement) is the sum of three random variables: time the fluorophore spend in the excited state; IRF time; and the number to pulses over which the fluorophore stays excited (also see eq. 12 in the main text). Consequently, the likelihood of a recorded photon arrival time, $\Delta t_k^i$, is given by the convolution of the probability distribution of these random variables as follows

$$P\left(\Delta t_k^i|\lambda_m\right) = \left[P\left(\Delta t_{\mathrm{IRF},k}^i\right|\tau_{\mathrm{IRF}},\sigma_{\mathrm{IRF}}^2) * P\left(\Delta t_{\mathrm{ext},k}^i|\lambda_m\right)\right] * P\left(n|N\right), \tag{S4}$$

where

$$P\left(n|N\right) = \mathrm{Categorical}_{0:N}\left([A_0, ..., A_N]\right) \tag{S5}$$
$$P\left(\Delta t_{\mathrm{IRF},k}^i\right|\tau_{\mathrm{IRF}},\sigma_{\mathrm{IRF}}^2) = \mathrm{Normal}(\Delta t_{\mathrm{IRF},k}^i;\tau_{\mathrm{IRF}},\sigma_{\mathrm{IRF}}^2) \tag{S6}$$
$$P\left(\Delta t_{\mathrm{ext},k}^i|\lambda_m\right) = \mathrm{Exponential}(\Delta t_{\mathrm{ext},k}^i;\lambda_m). \tag{S7}$$
$$\tag{S8}$$

Here, $\tau_{\mathrm{IRF}}, \sigma_{\mathrm{IRF}}^2, m$, $\lambda_m$ and $N$, respectively, denote the mean and variance of the IRF normal distribution, indicating the photon from the $k$th pulse within the $i$th pixel is from what species, the inverse of fluorophore lifetime, and the maximum number of pulses to be considered. Moreover, $A_n$ represent the probability of the photon originating from the $n$th previous pulse derived in [1]. Calculating the convolutions, we obtain the following likelihood model for the photon arrival time [1]

$$P\left(\Delta t_k^i|\lambda_m\right) = \left[\sum_{n=0}^{N}\mathrm{erfc}\left(\frac{\tau_{\mathrm{IRF}} - \Delta t_k^i - nT + \lambda_m\sigma_{\mathrm{IRF}}^2}{\sigma_{\mathrm{IRF}}\sqrt{2}}\right)\right.$$
$$\left.\times\ \frac{\lambda_m}{2}\exp\left(\frac{\lambda_m}{2}\left(2(\tau_{\mathrm{IRF}} - \Delta t_k^i - nT) + \lambda_m\sigma_{\mathrm{IRF}}^2\right)\right)\right]. \tag{S9}$$

Here, we continue by considering the probability $\pi_m^i$ of detecting a photon from the $m$th species in the $i$th pixel given by eq. 4 in the main text. Therefore, the species $m$ giving rise to a detected photon from a non-empty pulse can be sampled from a categorical distribution as follows

$$m|\Lambda_{1:M}, b_{1:M} \sim \mathrm{Categorical}_{1:M}\left(\pi_1^i, ..., \pi_M^i\right). \tag{S10}$$

Now, we can use eq. S10 to marginalize fluorophore species in the likelihood eq. S9 as follows

$$\begin{aligned}
P\left(\Delta t_k^i|\vartheta\right) &= \sum_{m=1}^{M} P\left(m|\Lambda_{1:M}, b_{1:M}\right) P\left(\Delta t_k^i|\lambda_m\right) \\
&= \left[\sum_{m=1}^{M}\pi_m^i P\left(\Delta t_k^i|\lambda_m\right)\right] \\
&= \left[\sum_{m=1}^{M}\pi_m^i\sum_{n=0}^{N} P(\Delta t_k^i + nT|\lambda_m)P(n|N)\right] \\
&= \left[\sum_{m=1}^{M}\pi_m^i\sum_{n=0}^{N}\mathrm{erfc}\left(\frac{\tau_{\mathrm{IRF}} - \Delta t_k^i - nT + \lambda_m\sigma_{\mathrm{IRF}}^2}{\sigma_{\mathrm{IRF}}\sqrt{2}}\right)\right. \\
&\left.\times\ \frac{\lambda_m}{2}\exp\left(\frac{\lambda_m}{2}\left(2(\tau_{\mathrm{IRF}} - \Delta t_k^i - nT) + \lambda_m\sigma_{\mathrm{IRF}}^2\right)\right)\right].
\end{aligned} \tag{S11}$$

The resulting marginal likelihood above does not depend on the fluorophore species $m$ and we employ it to develop our framework.

#### Supplementary Note 1.2: Priors and Posterior

After obtaining the likelihood, we proceed to build the object of most interest the posterior. The posterior is proportional to the product of the likelihood eq. S1 and prior distributions over the unknown parameters $\vartheta$. The entire model including the priors are summarized in the following. The posterior is given as

$$P\left(\vartheta|\overline{\overline{W}}, \overline{\overline{\Delta t}}\right) \propto P\left(\overline{\overline{\Delta t}}|\vartheta, \overline{\overline{W}}\right) P\left(\overline{\overline{W}}|\vartheta\right) P\left(\tau_{1:M}\right) P\left(\Lambda_{1:M}|\nu_{1:M}, b_{1:M}\right) P\left(\nu_{1:M}\right) P\left(b_{1:M}\right), \tag{S12}$$

where

$$
\begin{aligned}
W_k^i &\sim \text{Bernoulli}\left(1-\pi_0^i\right) && \text{(S13)}\\
\lambda_m &\sim \text{Gamma}\left(\alpha_\lambda, \beta_\lambda\right) && \text{(S14)}\\
\Lambda_m &\sim \mathbf{GP}\left(\nu_m, \boldsymbol{K}\left(\vec{X}, \vec{X}'\right)\right) && \text{(S15)}\\
\nu_m &\sim \text{Normal}\left(0, \sigma_\nu\right) && \text{(S16)}\\
b_m &\sim \text{Bernoulli}\left(b_m; \frac{1}{1+\frac{M-1}{\gamma}}\right), && \text{(S17)}
\end{aligned}
$$

where $\gamma$ is the expected number of species, $\alpha_\lambda, \beta_\lambda$ and $\sigma_\nu$ are hyper-parameters of gamma prior and sigma of prior on $\nu_m$. Moreover, we used $\lambda_m = 1/\tau_m$ and employ this parameter instead of $\tau_m$ from hereon. $\alpha_\lambda, \beta_\lambda$ and $\sigma_\nu$ are hyper-parameters of gamma prior and sigma of prior on $\nu_m$. The covariance matrix of the GP is given by

$$
\boldsymbol{K}\left(\vec{X}, \vec{X}'\right) = \sigma_{\text{GP}}^2 \exp\left[-\frac{1}{2}\left(\frac{\vec{X}-\vec{X}'}{L}\right)^2\right], \tag{S18}
$$

where $\sigma_{\text{GP}}$ and $L$ are positive parameters. Finally, we have the likelihood

$$
\begin{aligned}
P\left(\Delta t_k^i \middle| \vartheta\right) = &\left[\sum_{m=1}^{M} \pi_m^i \sum_{n=0}^{N} \text{erfc}\left(\frac{\tau_{\text{IRF}} - \Delta t_k^i - nT + \lambda_m \sigma_{\text{IRF}}^2}{\sigma_{\text{IRF}}\sqrt{2}}\right)\right.\\
&\left.\times \frac{\lambda_m}{2} \exp\left(\frac{\lambda_m}{2}\left(2(\tau_{\text{IRF}} - \Delta t_k^i - nT) + \lambda_m \sigma_{\text{IRF}}^2\right)\right)\right]^{W_k^i}.
\end{aligned} \tag{S19}
$$

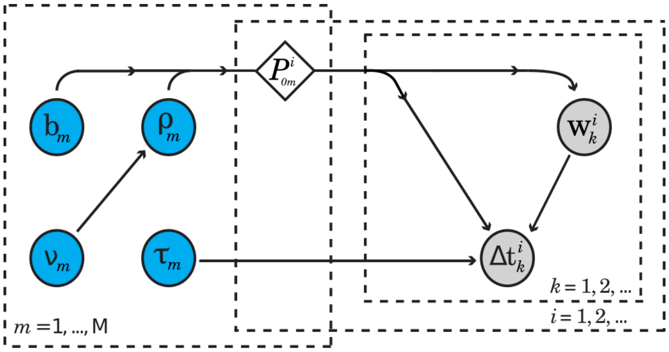


Fig. S9: Graphical Model. Grey and blue circles and the diamond, respectively, indicate observations, unknown parameters and variables that are detrministically calculated form other parameters. $k, i$ and $m$, respectively, count pulses, confocal regions and species. $W_k^i$ and $\Delta t_k^i$ reports if a pulse was empty or not and the micro-time of a photon from the $k$th pulse in $i$th confocal region, in turn. $P_{0m}^i$ represents the probability of the species $m$ not being excited during the $i$th pulse. $\rho_m$, $\tau_m$ and $b_m$ are, respectively, the concentration, lifetime and associated load of the $m$th species. $\nu_m$ is the parameter of the GP prior. Further, to facilitate computations, we need to work within a limited but large number of lifetimes $M$.

#### Supplementary Note 2: Model Inference

After deriving the posterior, in this section, we will provide the inverse strategy to infer the unknown parameters. Each parameter is deduced using the corresponding target distribution by sampling it either directly or using Metropolis-Hasting (MH) scheme.

#### Supplementary Note 2.1: Sampling $\lambda_m$

The target distribution of $\lambda_{1:M}$ is given by

$$\lambda_{1:M} \sim P\left(\lambda_{1:M}|\overline{\overline{\Delta t}}, \overline{\overline{W}}, \Lambda_{1:M}, \nu_{1:M}, b_{1:M}\right) \tag{S20}$$

$$\propto P\left(\overline{\overline{\Delta t}}|\vartheta\right) P\left(\lambda_{1:M}\right) \tag{S21}$$

$$= \left[\prod_i \prod_k P\left(\Delta t_k^i|\vartheta\right)\right] \prod_m P(\lambda_m) \tag{S22}$$

$$= \left[\prod_i \prod_k \left[\sum_{m=1}^{M} \pi_m^i \sum_{n=0}^{N} \frac{\lambda_m}{2} \exp\left(\frac{\lambda_m}{2}\left(2(\tau_{\text{IRF}} - \Delta t_k^i - nT) + \lambda_m \sigma_{\text{IRF}}^2\right)\right) \times \text{erfc}\left(\frac{\tau_{\text{IRF}} - \Delta t_k^i - nT + \lambda_m \sigma_{\text{IRF}}^2}{\sigma_{\text{IRF}}\sqrt{2}}\right)\right]^{W_k^i}\right] \left[\prod_m \text{Gamma}(\lambda_m; \alpha_\lambda, \beta_\lambda)\right], \tag{S23}$$

where the superscript $i$ is the index of pixels. The above expression does not have a closed form and therefore, Metropolis-Hastings (MH) algorithm [2,3] is used to sample $\lambda_{1:M}$. The proposed lifetimes, $\lambda_m^{\text{prop}}$, are taken from a gamma distribution

$$\lambda_{1:M} \sim \text{Gamma}\left(\alpha_\lambda^{\text{prop}}, \frac{\lambda_{1:M}^{\text{old}}}{\alpha_\lambda^{\text{prop}}}\right), \tag{S24}$$

and the MH acceptance ratio is given by

$$A = \frac{P\left(\lambda_{1:M}^{\text{prop}}|\overline{\overline{\Delta t}}, \overline{\overline{W}}\right) \text{Gamma}\left(\lambda_{1:M}^{\text{old}}; \alpha_\lambda^{\text{prop}}, \frac{\lambda_{1:M}^{\text{prop}}}{\alpha_\lambda^{\text{prop}}}\right)}{P\left(\lambda_{1:M}^{\text{old}}|\overline{\overline{\Delta t}}, \overline{\overline{W}}\right) \text{Gamma}\left(\lambda_{1:M}^{\text{prop}}; \alpha_\lambda^{\text{prop}}, \frac{\lambda_{1:M}^{\text{old}}}{\alpha_\lambda^{\text{prop}}}\right)}. \tag{S25}$$

#### Supplementary Note 2.2: Sampling $\Lambda_m$

The target distribution of $\Lambda_{1:M}$ is given by

$$\Lambda_{1:M} \sim P\left(\Lambda_{1:M}|\overline{\overline{\Delta t}}, \overline{\overline{W}}, \lambda_{1:M}, \nu_{1:M}, b_{1:M}\right) \tag{S26}$$

$$\propto P\left(\overline{\overline{\Delta t}}, \overline{\overline{W}}|\lambda_{1:M}, \Lambda_{1:M}, b_{1:M}\right) P\left(\Lambda_{1:M}|\nu_{1:M}, b_{1:M}\right), \tag{S27}$$

where

$$P\left(\overline{\overline{\Delta t}}, \overline{\overline{W}}|\lambda_{1:M}, \Lambda_{1:M}, b_{1:M}\right) = P\left(\overline{\overline{\Delta t}}|\vartheta\right) P\left(\overline{\overline{W}}|\Lambda_{1:M}, b_{1:M}\right) = \left[\prod_i \prod_k P\left(\Delta t_k^i|\vartheta\right) \left(1 - \pi_0^i\right)^{W_k^i} \left(\pi_0^i\right)^{1-W_k^i}\right], \tag{S28}$$

where $\pi_0^i$ is given by eq. 6 in the main text. We take GP priors on the species concentrations, given in eq. S15.

While GP priors allow negative values, molecular concentrations and thus lifetime maps ($\Lambda_{1:M}$) are positive quantities and thus we use the following substitution to assure that the proposed values are always non-negative

$$\Lambda_m = \exp(\chi_m) \tag{S29}$$

where $\chi_m$ takes both negative and positive values. We learn $\chi_{1:M}$ employing the GP method and then find the concentrations using eq. S29. The likelihood of $\chi_{1:M}$, $P\left(\overline{\overline{\Delta t}}, \overline{\overline{W}}|\lambda_{1:M}, \chi_{1:M}, b_{1:M}\right)$, is obtained by replacing $\Lambda_{1:M}$ with $\exp(\chi_{1:M})$ in eq. S28. The resulting likelihood is non-conjugate to the GP prior and hence the posterior does not have a closed form. As such, the posterior cannot be directly sampled and we need to adopt either the MH or the elliptical slice sampling [4] techniques to make inference about $\chi_{1:M}$. After conducting some test experiments with synthetic data, we decided to opt for the elliptical slice sampling with the following steps:

**Step 1:**
Select a random lifetime $m$

**Step 2:**
Select a random number $u \in [0,\, 1]$

**Step 3:**
Set $\theta_{\min} = 0$ and $\theta_{\max} = 2\pi$ and
select a random number $\theta \in [\theta_{\min},\, \theta_{\max}]$

**Step 4:**
Select a random set of parameters from the prior on the $m$th lifetime $\chi_m^* \sim \mathbf{GP}(\nu_m, \mathbf{K})$

**Step 5:**
Propose a new lifetime map as follows

$$\chi'_m = \chi_m \cos\theta + \chi_m^* \sin\theta \tag{S30}$$

**Step 6:** Calculate the ratio of the likelihoods for the current and proposed maps

$$A = \frac{P\left(\overline{\overline{\Delta t}}, \overline{\overline{W}} | \lambda_{1:M}, \chi'_{1:M}, b_{1:M}\right)}{P\left(\overline{\overline{\Delta t}}, \overline{\overline{W}} | \lambda_{1:M}, \chi_{1:M}, b_{1:M}\right)}, \tag{S31}$$

where the prime denotes the set of $\chi$-maps where the $m$th map is given by the map generated above while the rest stay the same.

**Step 7:**
Accept the proposal if $A > u$. Otherwise, set
$\theta_{\min} = \theta$ if $\theta < 0$, else
$\theta_{\max} = \theta$ if $\theta > 0$, and
Select at random $\theta \in [\theta_{\min},\, \theta_{\max}]$

**Step 8**
Go to step 4 and repeat until accepting a proposal.

#### Supplementary Note 2.3: sampling $\nu_m$

We take a normal distribution as prior on the mean of GP priors, $\nu_{1:M}$, and learn them. The full conditional posterior of $\nu_{1:M}$ is

$$\begin{aligned} P\left(\nu_{1:M} | \chi_{1:M}\right) &\propto P\left(\chi_{1:M} | \nu_{1:M}\right) P\left(\nu_{1:M}\right) \\ &= \mathbf{GP}\left(\chi_{1:M}; \nu_{1:M}, \boldsymbol{K}\right) \boldsymbol{\mathcal{N}}\left(\nu_{1:M}, 0, \sigma_\nu^2\right) \end{aligned} \tag{S32}$$

where $\sigma_\nu^2$ is the variance of the normal prior. The values of $\nu_{1:M}$ are modified using the MH method where the acceptance ratio is

$$A = \frac{\mathbf{GP}\left(\chi_{1:M}; \nu'_{1:M}, \boldsymbol{K}\right) \boldsymbol{\mathcal{N}}\left(\nu'_{1:M}, 0, \sigma_\nu^2\right)}{\mathbf{GP}\left(\chi_{1:M}; \nu_{1:M}, \boldsymbol{K}\right) \boldsymbol{\mathcal{N}}\left(\nu_{1:M}, 0, \sigma_\nu^2\right)}, \tag{S33}$$

where prime indicate the proposed values obtained using a random walk.

#### Supplementary Note 2.4: Sampling $b_m$

The full target distribution of binary weights is

$$
\begin{aligned}
b_{1:M} &\sim P\left(b_{1:M}\middle|\overline{\overline{\Delta t}}, \overline{\overline{W}}, \lambda_{1:M}, \Lambda_{1:M}, \nu_{1:M}\right) \\
&\propto P\left(\overline{\overline{\Delta t}}|\vartheta\right) P\left(\overline{\overline{W}}|\Lambda_{1:M}, b_{1:M}\right) P\left(b_{1:M}\right) \\
&= \left[\prod_i \prod_k \left[\sum_{m=1}^{M} \pi_m^i \sum_{n=0}^{N} \frac{\lambda_m}{2} \exp\left(\frac{\lambda_m}{2}\left(2(\tau_{\text{IRF}} - \Delta t_k^i - nT) + \lambda_m \sigma_{\text{IRF}}^2\right)\right)\right.\right. \\
&\times \left.\left. \text{erfc}\left(\frac{\tau_{\text{IRF}} - \Delta t_k^i - nT + \lambda_m \sigma_{\text{IRF}}^2}{\sigma_{\text{IRF}}\sqrt{2}}\right)\right]^{W_k^i} \left(1-\pi_0^i\right)^{W_k^i} \left(\pi_0^i\right)^{1-W_k^i}\right] \\
&\times \left[\prod_m \text{Bernoulli}\left(\mathrm{b_m}; \frac{1}{1+\frac{\mathrm{M}-1}{\gamma}}\right)\right],
\end{aligned}
\tag{S34}
$$

where the likelihood and prior are given by eq. S28 and eq. S17, respectively. Here, since an infinite number of lifetimes is computationally formidable, we rather use a large but limited number of lifetimes $M$ within the model. Moreover, the prior parameter $1/\left(1+\frac{M-1}{\gamma}\right)$ is obtained by marginalization over the beta-Bernoulli process [5–7].

To sample the loads associated to the $M$ present lifetimes in the model, we randomly pick two of them to update while the remaining loads stay fixed. Now, there are four possibilities associated to the two selected loads

$$
\begin{aligned}
\overline{B}_1 &= [0, 0] \\
\overline{B}_2 &= [1, 0] \\
\overline{B}_3 &= [0, 1] \\
\overline{B}_4 &= [1, 1],
\end{aligned}
\tag{S35}
$$

where $\overline{B}_l$ represents the $l$th possible combination of loads. The loads can then be directly sampled from a categorical distribution as follows

$$
b_{1:M} \sim \text{Categorical}_{1:4}\left(\frac{P\left(\overline{B}_1|...\right)}{G}, ..., \frac{P\left(\overline{B}_4|...\right)}{G}\right)
\tag{S36}
$$

where $P\left(\overline{B}_l|...\right)$ denotes the posterior given in eq. S34 for the case where the two selected loads take the $l$th values and

$$
G = \sum_{l=1}^{4} P\left(\overline{B}_l|...\right).
\tag{S37}
$$

#### Supplementary Note 3: Table of Parameter Values used for Data Analysis

| Parameter | $\alpha_\lambda$ | $\beta_\lambda$ | $\alpha_\lambda^{prop}$ | $\sigma_{\mathrm{GP}}$ | $L$ | $\sigma_\nu$ |
|---|---|---|---|---|---|---|
| Unit | - | ns | - | $\mu$m | $\mu$m | $\mu$m |
| Fig. 2 | 1 | 5 | 2,000 | 1 | 1 | 50 |
| Fig. 3 | 1 | 5 | 2,000 | 1 | 0.4 | 50 |
| Fig. 4 | 1 | 5 | 2,000 | 1 | 0.4 | 50 |
| Fig. 5 | 1 | 5 | 2,000 | 1 | 0.5 | 50 |
| SI Fig. 1 | 1 | 5 | 2,000 | 1 | 1 | 50 |
| SI Fig. 2-6 | 1 | 5 | 2,000 | 1 | 0.4 | 50 |
| SI Fig. 7 | 1 | 5 | 2,000 | 1 | 0.5 | 50 |

Table S1: Table of the the parameter values used for the analysis of the data.

### References


[1] Mohamadreza Fazel, Sina Jazani, Lorenzo Scipioni, Alexander Vallmitjana, Enrico Gratton, Michelle A Digman, and Steve Pressé. High resolution fluorescence lifetime maps from minimal photon counts. *ACS photonics*, 9(3):1015–1025, 2022.

[2] Nicholas Metropolis, Arianna W Rosenbluth, Marshall N Rosenbluth, Augusta H Teller, and Edward Teller. Equation of state calculations by fast computing machines. *The journal of chemical physics*, 21(6):1087–1092, 1953.

[3] W Keith Hastings. Monte carlo sampling methods using markov chains and their applications. *Biometrika*, 57:97–109, 1970.

[4] Iain Murray, Ryan Adams, and David MacKay. Elliptical slice sampling. In *Proceedings of the thirteenth international conference on artificial intelligence and statistics*, pages 541–548. JMLR Workshop and Conference Proceedings, 2010.

[5] John Paisley and Lawrence Carin. Nonparametric factor analysis with beta process priors. In *Proceedings of the 26th annual international conference on machine learning*, pages 777–784, 2009.

[6] Luai Al Labadi and Mahmoud Zarepour. On approximations of the beta process in latent feature models: Point processes approach. *Sankhya A*, 80(1):59–79, 2018.

[7] Sina Jazani, Ioannis Sgouralis, Omer M Shafraz, Marcia Levitus, Sanjeevi Sivasankar, and Steve Pressé. An alternative framework for fluorescence correlation spectroscopy. *Nature communications*, 10(1):1–10, 2019.
